# Shape stabilization in budding intestinal organoids by geometrical ratcheting

**DOI:** 10.1101/2025.04.02.646749

**Authors:** Oliver M. Drozdowski, Kim E. Boonekamp, Ulrike Engel, Michael Boutros, Ulrich S. Schwarz

**Affiliations:** BioQuant, Heidelberg University, 69120 Heidelberg, Germany; Institute for Theoretical Physics, Heidelberg University, 69120 Heidelberg, Germany; Max Planck School Matter to Life, 69120 Heidelberg, Germany; German Cancer Research Center (DKFZ), Division Signaling and Functional Genomics and Medical Faculty Heidelberg, Heidelberg University, 69120 Heidelberg, Germany; Nikon Imaging Center and Centre for Organismal Studies, Heidelberg University, 69120 Heidelberg, Germany

## Abstract

The intestinal epithelium in vertebrates has a characteristic arrangement of protruding villi and receding crypts, which are generated and stabilized by selforganized cell forces. Intestinal organoids recapitulate this development and can be used to shed light on the underlying mechanical processes. Here we combine advanced image processing and the bubbly vertex model for epithelial cell shape to achieve a fully three-dimensional reconstruction of cell shapes, forces and tissue geometries during the development of mouse intestinal organoids. We show that the transition from spherical to budded morphologies is caused by a global increase in tension at the inner side and a decrease in luminal pressure, which however are not maintained after budding, suggesting ratchet-like non-reversibility. Through computer simulations of a vertex model for growing tissue with T1-transitions and comparison with the experimental data, we show that this ratchet is likely of geometrical origin. We further find that the system is mechanically stabilized on the outer side by increases in line tensions and reorganization of the actomyosin network, stabilizing a polygonal organization that also seems to enforce the geometrical ratchet. Our approach demonstrates how one can achieve a complete mechanical analysis of a complex tissue-like system.

## Introduction

To facilitate uptake of nutrients, the surface area in the intestine is increased by the convoluted shapes generated by the alternating arrangements of invaginations and protrusions, called crypts and villi, respectively [1]. Because cells in the intestine are mechanically and chemically strongly challenged by their environment, they are continuously expelled from the villi regions, with the typical turnover time of the entire intestinal epithelium being 3-5 days [2, 3]. In order for the intestine to adopt its characteristic shape, to generate the continuous flow of cells from the crypts to the villi, and to expel cells toward the villus tips, physical forces have to be generated. However, to date no method exists to measure such forces *in situ* and thus it is not clear how the overall system achieves mechanical stability.

Over the last decade, intestinal organoids, which are grown from a few stem cells into mini-versions of this organ, have emerged as a powerful model system to address fundamental questions on the intestine [4, 5]. In particular, it has been shown that intestinal organoids undergo a morphological transition from a spherical to a budded shape, recapitulating key developmental characteristics of the gut, but also pointing to morphological transformations during cancer [6, 7]. Intestinal organoids have been used for drug screening and disease modelling [8] and are very appealing model systems to study tissue mechanics [9].

It is well established that the main driver for cell shape changes in epithelial tissue like the intestine is the actomyosin system, generating forces mainly in the form of cortical surface tension [3, 10]. Asymmetries in tensions between inner (apical) and outer (basal) interfaces have been shown to be involved in the onset of intestinal organoid budding [6, 7]. Cell extrusion, where single cells are pushed out of the monolayer, has been linked to both tissue-scale tension and force generation in neighboring cells [11, 12]. Typical signatures of surface tension in cellular systems are spherical shapes at free surfaces, invaginated arcs between regions of strong adhesion, and polyhedral shapes for close packing [13]. Based on the concept of surface tension, the vertex model has emerged as the standard mathematical model to describe the structure of epithelia in three dimensions [6, 7, 14–17]. However, the current focus on surface tensions neglects other potential contributions to mechanical stability, including luminal pressure, intracellular structures like the nucleus, and line tensions.

In order to understand how epithelia like the intestine control shape changes and mechanical stability, one can measure cell forces by laser cutting [6, 18], embedded force probes [19–21], or on soft elastic substrates [22]. Subjecting epithelia to deformations also probes their mechanics [23–26]. A non-invasive, but challenging alternative is force inference, which has been successfully implemented for two-dimensional (2D) epithelial sheets [27–32], but has not been achieved yet for the fully three-dimensional (3D) structure of epithelia. Previous approaches in 3D have primarily focused on small animal and human embryos [33–36] or more recently larger embryos [37, 38], but intestinal organoids contain more intracellular structure than embryos, thus image quality and force inference are more challenging. In particular, they have a stronger cytoskeleton and thus one expects not only surface tensions, but other mechanical processes to contribute, including previously under-appreciated line tensions. Another important aspect are viscoelastic relaxation processes, which also are strongly determined by the actomyosin system [23].

To investigate the role of cell forces for organoid tissue mechanics, we have developed a fully three-dimensional geometry reconstruction and force inference pipeline for intestinal organoids, overcoming the existing challenges of image quality and mechanical force balance in 3D. We find that inner (apical) tensions are globally increased before budding, then trigger bud formation, but are not used to stabilize existing buds, suggesting a ratchet-like irreversibility of budding. We also show that luminal pressure facilitates the budding process and nuclear shape is not affected much by the morphological transition. Through inference of organoid topology and computer simulations, we investigate tissue-scale dynamics and show that a geometrical ratchet resulting from the interplay of growth and fluidization stabilizes organoid shape. Lastly, we demonstrate the existence of outer (basal) line tensions, which are a signature of additionally generated forces that do not correspond to surface tensions and which result from a cytoskeletal reorganization into a more polygonal structure. This stabilizes the topological structure of the organoid and thus also seems to be an essential element of shape stabilization.

## Results

### Cell geometries and forces can be fully reconstructed in 3D

Mouse small intestinal organoids were grown in extracellular matrix (basement membrane extract) after mechanical splitting of budded organoids, to release crypt regions with the capacity to form new organoids (see methods). Released crypts close into symmetrical spherical shells after 5 h, undergo symmetry breaking into an elongated shape after 48 h and on average display a complex budded morphology after 53 h (Fig. 1a). After matrix removal, excluding effects from tissue-environment interactions [39–43], the organoids were fixed and stained and then using fluorescence microscopy, 3D imaging data were acquired (Fig. 1b). In order to determine the cell-cell interfaces, the cells in the organoids were segmented voxel-wise [44–46] (Fig. 1c, Supplementary Video 1). For the reconstruction we developed a procedure based on non-parametric statistics of point clouds [34], from which we obtain the normal vectors of the interfaces in a spatially resolved manner (Fig. 1d,e).

**Fig. 1.**
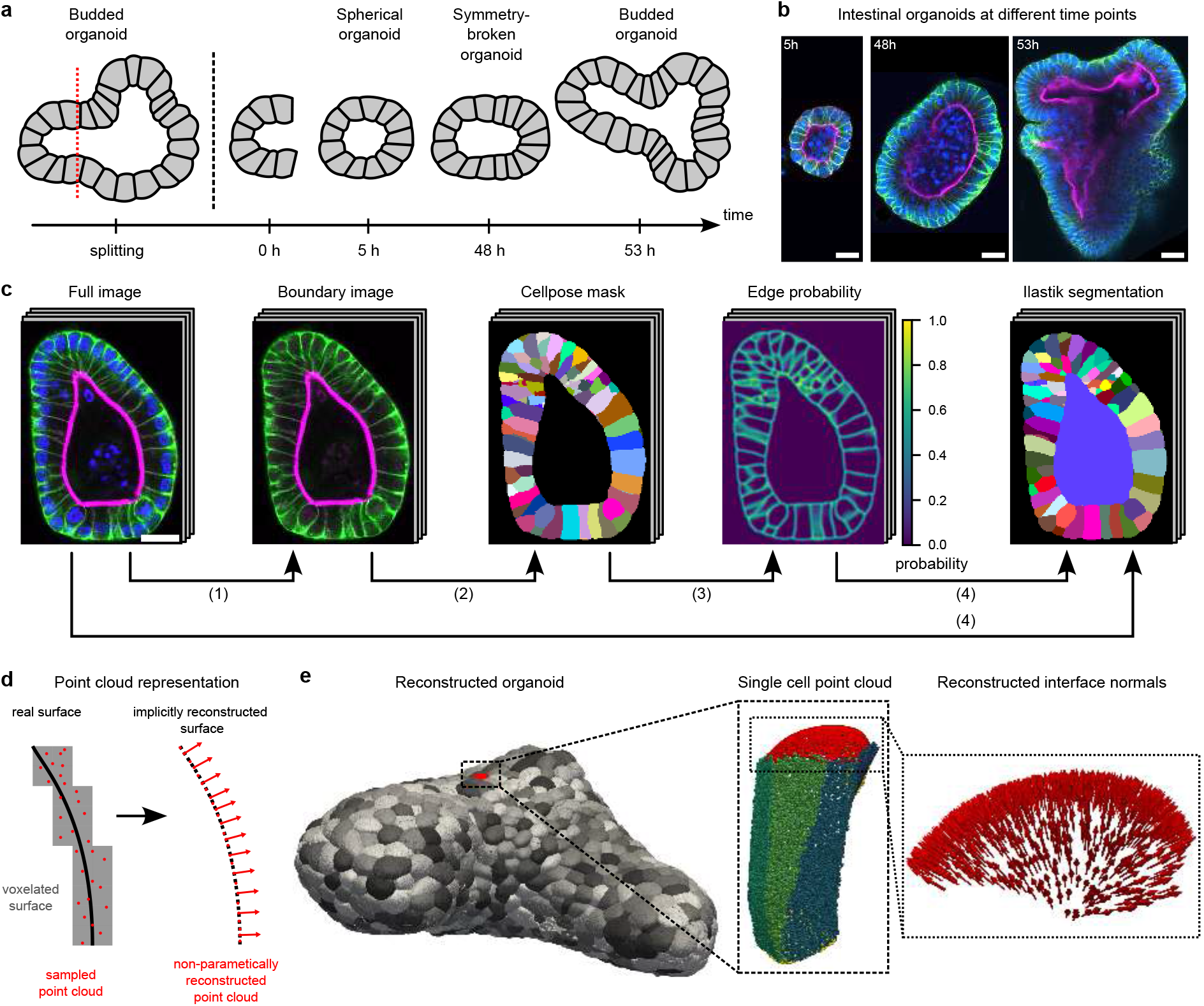
Segmentation and three-dimensional geometry reconstruction of mouse small intestine organoids. **a**, During passaging of the mouse small intestinal organoids, the crypts are dissociated from the fully grown organoid structures. The crypt region containing stem cells subsequently serves as starting point for the development of another organoid. 5 hours post-splitting the organoids close and form a spherical shell. At 48 hours post-splitting the spherical symmetry is broken, the organoid bulges and starts to form new crypt regions. At 53 hours post-splitting the organoids display a complex budded shape. **b**, Example microscopy images of organoids at different experimental time points. F-actin is in magenta, *β*-catenin in green and nuclei in blue. **c**, Segmentation pipeline of 3D microscopy data (and similarly for simulated image stacks) using Cellpose and Ilastik. See methods for a detailed explanation. Probability map of cell edges is color coded by probability and in segmentations individual cells are arbitrarily colored. Scale bars in **b, c** are 20 *µ*m. **d**, From the segmentation a voxelated representation of interfaces can be determined, from which in turn a point cloud based representation is created. This point cloud is used to non-parametrically infer the underlying surface with the corresponding normal vectors (see methods for details). **e**, Example of reconstruction results for a budded organoid on different scales. The organoid consists of several hundreds cells. One cell consists of typically eight different interfaces, which are represented as point clouds. For each point in the surface, our algorithm also constructs a normal vector, which is important for angle determination.

Based on the successful geometry reconstruction, we then implemented a fully 3D force inference pipeline. In addition to reconstruction of surface tension, which is at the center of 2D force inference, 3D force inference also allows to reconstruct line tensions, and thus to obtain complete force balance at the lines and points at which different cells connect (Fig. 2a). In order to validate our method and to estimate the spatial resolution, we use computer simulations of virtual organoids with the bubbly vertex model (BVM) [47–50], which in contrast to the traditional vertex model allows for curved interfaces (Fig. 2b). The procedure starts by using the reconstructed normal vectors to calculate the dihedral angles of intersecting interfaces at tri-interfacial junctions (Fig. 2c). Force balance of three interfaces with (relative) tensions *γ*_*i*_ (*i* = 1, 2, 3) can be formulated via

**Fig. 2.**
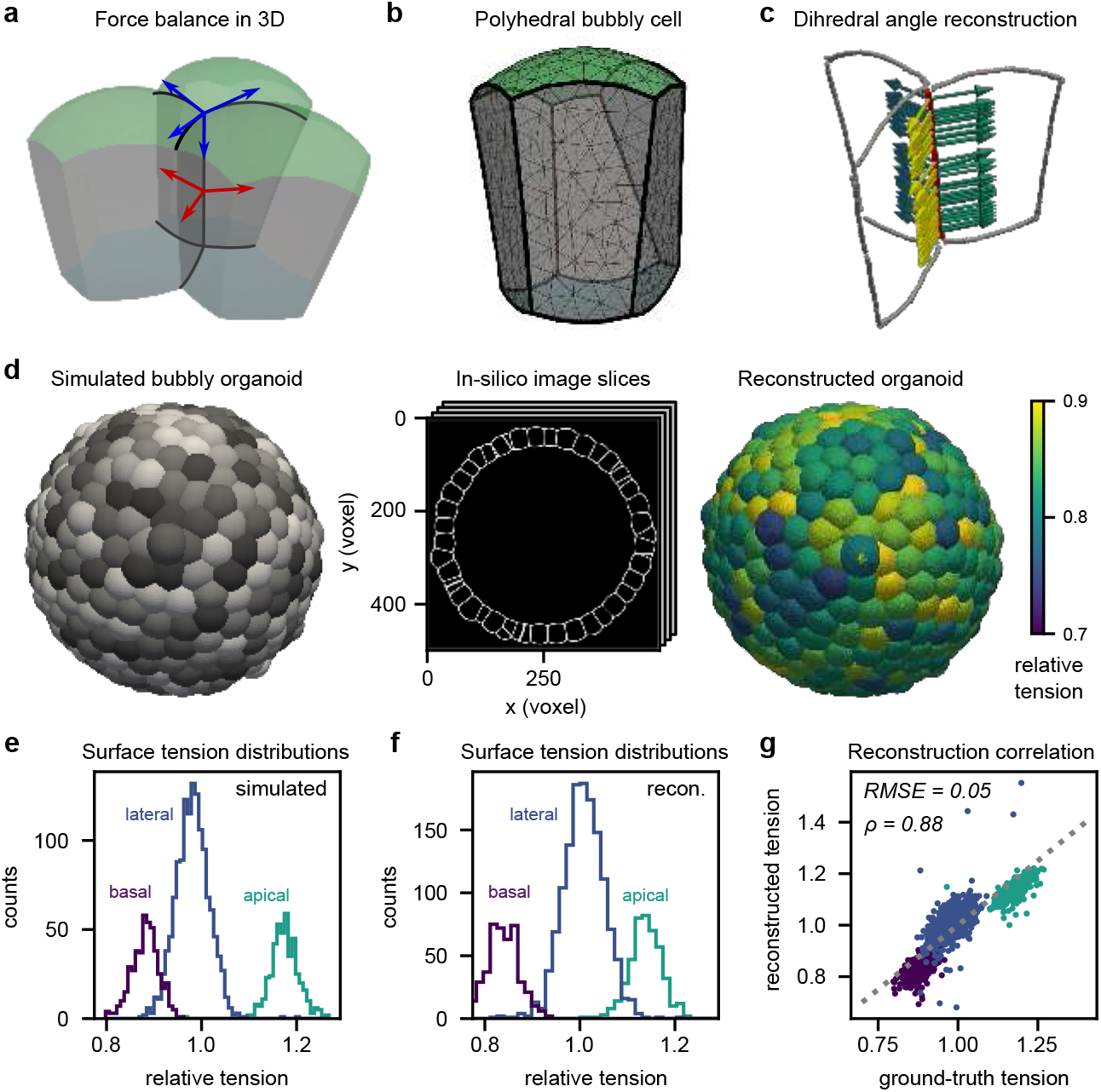
In silico validation of the three-dimensional force inference. **a**, Force balance at tri-interfacial junctions (red) and at vertices (blue) enable the inference of forces and thus surface and line tensions in 3D. **b**, Cells in organoids display a polyhedral shape with curved interfaces, which can be simulated in silico with the bubbly vertex model. **c**, The dihedral angles between the interfaces from an organoid (here in silico) are determined from the reconstructed surface tangents (arrows) along a tri-interfacial junction (red). **d**, With the bubbly vertex model an organoid is simulated, image stacks are created with a similar resolution as the experimental data, and the reconstruction and force inference pipeline is applied. **e,f**, Distributions of different interfaces by type in (**e**) ground truth simulations and (**f**) inferred reconstructions. g, Correlation of ground truth simulated surface tensions and inferred relative surface tensions, with root mean squared error (RMSE) and Pearson correlation coefficient *ρ*.

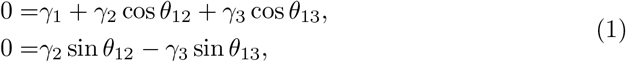

with angles *θ*_*ij*_ between interfaces *i* and *j*. This approach has been demonstrated before to be suitable for 3D force inference [34, 37]. Equation (1) defines a linear problem for the surface tensions, which we solve in a least squared sense. Because these equations can define only relative surface tensions, we fix the reference surface tension to the value 1, using 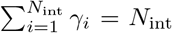 for *N*_int_ interfaces in total. From published work, we know that a typical value for the reference surface tension would be 1 *nN/µm* =1 *mN/m* [6].

We computationally validated our force inference method in a two-cell test system which can be solved analytically (extended data Fig. E1)—finding a typical error of force in the range of 5 percent (see Supplementary Notes 2 and 3 for details)— and in large systems with hundreds of cell simulated with the BVM (Fig. 2d). As the inference problem constitutes a noisy inverse problem, we used Thikonov regularization [51] in the least squared error estimation (see Supplementary Note 2). We found that the ground truth distribution of tensions (Fig. 2e) is very well reconstructed by our method (Fig. 2f), with a very strong correlation between the individual tensions (Fig. 2g, extended data Fig. E2).

### Budding is not maintained yet caused by apical tension

After successful benchmarking of our automated and completely 3D force inference pipeline, we applied it to experimental data. At 53 h post-splitting, we detected both budded and non-budded morphologies. We performed our force inference on these organoids, yielding a point-cloud representation of the organoid geometries with a value for the tension assigned to each of the basal, lateral and apical interfaces (Fig. 3a, Supplementary Video 2). Assessing the individual tensions of a representative organoid, we find skewed and broad distributions (Supplementary Figure S1). Occasional segmentation errors like seen in Fig. 3a can in general be neglected because of the large data sets for our statistical analysis (the number of interfaces is of the order of thousands). Organoids were gathered in different batches and from different mouse donors with batch-to-batch variability (Supplementary Figure S2), which we corrected for by performing a normalization of the apical tension deviation from the average value (see methods).

**Fig. 3.**
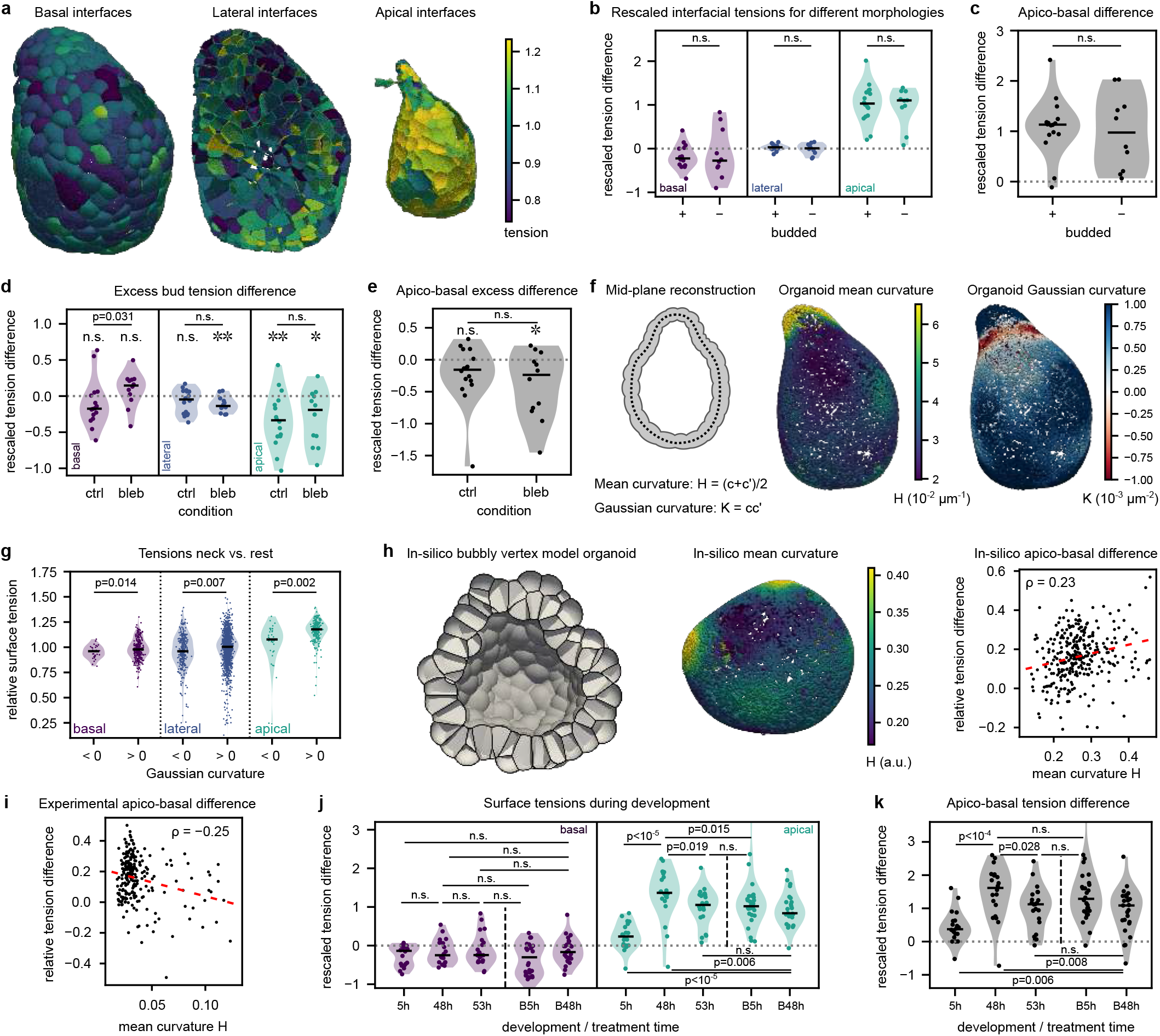
Force inference of surface tensions in budded and non-budded intestinal organoids suggests ratchet-like bud stabilization. **a**, Point-cloud reconstruction of organoid interfaces 53 h post-splitting, color-coded by relative surface tension. **b**, Tension differences from average tension by interface type, rescaled as to be normalized by mouse donor (see methods), for budded vs. non-budded morphologies. **c**, Rescaled single-cell apico-basal tension differences for budded vs. non-budded morphologies. **d**, Rescaled excess bud tension, which is the difference of the median rescaled tension (per organoid) in the bud and the rest by interface type for 53 h post splitting with (bleb) and without (ctrl) 5 h blebbistatin treatment. Negative values indicate tension in the bud is lower. **e**, Difference of apical and basal excess tensions per organoid. Negative values mean that the apico-basal difference is lower in the bud regions. **f**, Reconstructed mid-plane of the organoid with mean curvature *H* = (*c* + *c*^′^)*/*2 and Gaussian curvature *K* = *cc*^′^, where *c, c*^′^ denote the principle curvatures. **g**, Interfacial tensions by type in negative Gaussian curvature region (neck) and positive Gaussian curvature region (rest). As statistical test a two-sample two-sided t-test of independent samples with unequal variance was used. We find lower tensions in the neck region. **h**, Cross-section of budded simulated (in silico) organoid with corresponding mean curvature reconstruction and ground truth single-cell apico-basal tension differences as function of mid-plane curvatures. Red line is a linear fit, indicating the positive correlation with Pearson correlation coefficient *ρ* in the surface-tension based morphology. **i**, Single-cell apico-basal tension differences with corresponding correlations with mean curvature for experimental organoid. Red line is the linear fit, indicating the negative corelation in experimental data. **j**, Rescaled tension median distributions for organoids at different developmental time points with and without blebbistatin treatment. Interfacial tensions at 5 h, 48 h, and 53 h post-splitting and with 5 h blebbistatin treatment after 48 h (B5h) and with 48 h treatment after 5 h (B48h). **k**, Corresponding apico-basal tension difference medians. As statistical test a Mann-Whitney-U-test was used. Confidence levels ∗: *p* ≤ 0.05; ∗∗: *p* ≤ 0.01. Black lines in violin plots indicate medians. Color-scale boundaries adjusted to see sign change in Gaussian curvature (f) and to 5% and 95% quantiles in mean curvatures and tensions (**a,f,h**).

Our first main finding is that apical tension is higher than basal and lateral tensions in a statistically significant manner (Fig. 3b, Supplementary Figure S1). This agrees with the observation that the apical side is characterized by strong accumulation of actin (Fig. 1b). The single-cell apico-basal tension difference *γ*_**a**_ − *γ*_**b**_ shows a broad distribution which peaks at positive values, which effectively corresponds to a spontaneous curvature, consistent with the inward curvature (Supplementary Figure S1). Strikingly, we find that the rescaled tensions and apico-basal tension difference do not show any significant differences between budded and non-budded morphologies (Fig. 3b,c). This is surprising, because one might have expected that buds are maintained locally by higher apical tensions, and suggests that the higher apical tension is not required to maintain the budded morphology. To verify this conclusion, we determined the excess bud tensions, i.e. the increase of tensions within budded areas of organoids compared to the rest (cf. Supplementary Video 3). We found that apical (significantly) and basal (non-significantly) excess tensions are negative, suggesting that tensions are in fact higher outside the bud regions (Fig. 3d,e). Treatment with blebbistatin, an inhibitor of actomyosin contraction, for 5 hours before the 53 h time point leads to less and weaker budding. We find increases of the excess bud tensions basally, indicating the tensions generated outside the buds are driven by actomyosin contraction (Fig. 3d,e).

The successful reconstruction of all surface tensions allows us also to describe the whole organoid as a thin shell enclosing a lumen. To this end, we performed a pointcloud-based non-parametric reconstruction of the organoid mid-plane for a budded organoid, similarly to interface reconstruction (see methods and Supplementary Note 1), to obtain the tissue mid-plane and its mean and Gaussian curvatures with high spatial resolution (Fig. 3f, Supplementary Video 4). We find that the neck regions display lower tensions, in agreement to expected smaller spontaneous curvature (Fig. 3g). However, tissue curvature and the apico-basal tension difference, which correlate in the BVM, are anti-correlated in experiments (Fig. 3h,i, Supplementary Video 5), suggesting that surface tensions do not fully explain the observed tissue curvature, in agreement with our conclusion that buds are not maintained by apical contraction.

While the increase in apical tension seems not to be required to maintain budding, it is likely to be required to cause budding, as suggested by earlier studies [6, 7, 10]. To clarify this important point, we performed force inference at different stages of structure formation and found that indeed apico-basal tension differences are increased at the symmetry breaking stage around 48 h (Fig. 3j,k). Strikingly, they then decrease again, in agreement with our conclusion that they are not required for bud maintenance. We hypothesize that after budding, maintenance of shape is ensured by stabilizing the new geometry in a more permanent manner, and possibly without active actomyosin forces. Thus budding might work like an irreversible ratchet, as described before for other developing systems [52–54] and recently also for optic vesicle organoids [55]. We note that the budding process takes around 15 minutes [6, 10]; here we do not resolve such short periods of times, but reconstruct dynamical trends in apical tension over the whole developmental time.

In addition to actomyosin tensions, also nuclei have been proposed to contribute to the mechanobiology of cells [56–58] and have been used for stress inference in tissues [59]. To rule out that nuclear mechanics drive the budding process, we segmented nuclei and analyzed their volumes and shapes (Extended Data Fig. E3). We found that indeed nuclei do not contribute to the budding and do not show shape differences in our free organoids throughout development. This is in marked contrast to intestinal organoids in scaffolds, for which recently strong nuclear deformations with effects on signaling have been observed [60], suggesting that nuclei have a particularly important role for interactions with rigid environments [42]. For organoids grown in soft extracellular matrix as used here, such a coupling is unlikely.

Leveraging the Young-Laplace law, we can also infer luminal pressures (Extended Data Fig. E4), which we verified on in-silico data (Supplementary Note 3). For experimental data, we do not see systematic patterning in cellular pressures (Extended Data Fig. E4b,c). We see that the rescaled luminal pressure decreases significantly at the 48 h symmetry breaking time point (Extended Data Fig. E4d,e), which is consistent with previous observations of lumen shrinkage at the onset of budding [6]. The pressure increases after budding, however, which implies that it does not contribute to shape stabilization.

### Ratchet in local tissue geometry stabilizes organoid shape

To better understand which physical mechanisms might stabilize shape after budding, we now turn to the dynamic aspects of organoid growth. Intestinal organoids grow through cell division of stem cells in the buds into transit amplifying cells which differentiate upon further division [22]. To explore the dynamics of three-dimensional shape generation, we investigated growing organoids *in silico*. We considered the flat-interface limit of the bubbly vertex model, i.e. the classical vertex model (VM) [17] and introduced cell divisions. In addition we introduced T1-transitions, i.e. cell-cell rearrangements in the VM, which allows the tissue to adapt to altered stress like a fluid. We find that shells with small tissue fluidity (small T1-transition rate per cell) form buds akin to tip growth, which have been described before [61] (Fig. 4a, Supplementary Video 6) and are consistent with experimentally observed shapes of some of our organoids (Fig. 4b). However, due to the linear increase in cell number (Fig. 4c), the size increases and with a constant rate of T1-transitions per cell we find destabilization of the symmetric spherical configuration with time (Fig. 4d). For constant fluidity therefore cell division leads to morphological symmetry breaking. This is contrary to the case of constant size, where fluidity leads to more complex shapes [17]. This can be understood in a simple scaling argument: Assuming random cellular motion on the sphere, both the typical length scale of movement and the distances on the growing sphere scale with time as ∝ *t*^1*/*2^. If the shell is too large, the necessary distance that cells need to move to maintain spherical shapes increases and cannot be compensated anymore. We thus find a geometrical ratchet where fluidity cannot overcome growth beyond a typical size.

**Fig. 4.**
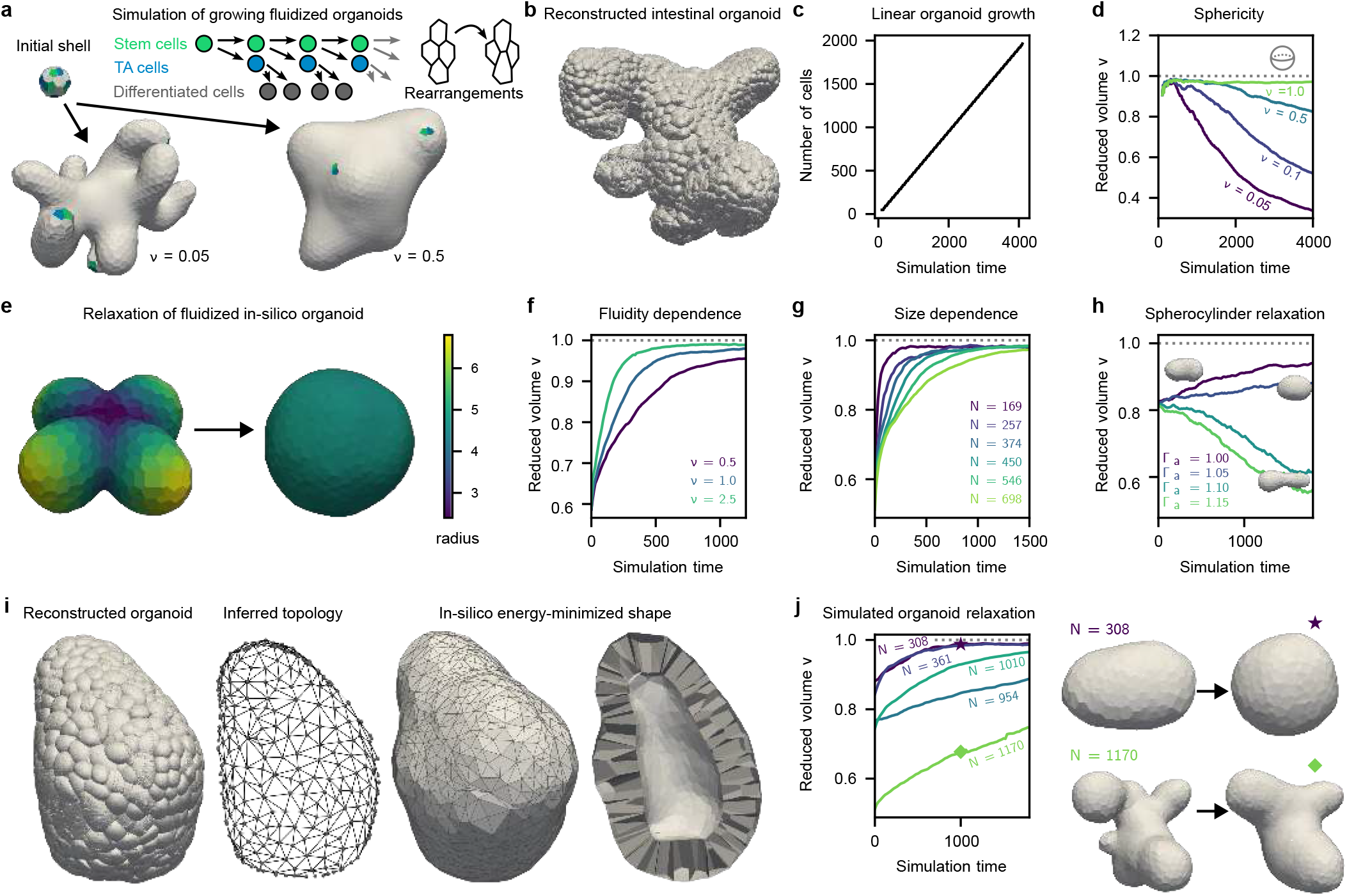
Geometrical ratchet stabilizes organoid shape. **a**, Flat-interface vertex model (VM) simulation of growing organoids—stem cells divide into transit amplifying (TA) cells, which divide into differentiated cells—with cell-cell rearrangements (T1 transitions) fluidizing the tissue. For larger T1 rate per cell *ν* (fluidity) the cell remains more spherical. **b**, Three-dimensional reconstruction of a complex intestinal organoid at 53 h. **c**, For the intestinal-like cell division pattern the number of cells grows linearly with simulation time. **d**, The reduced volume, 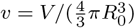 with the volume *V* and radius of a sphere corresponding to area *A*, 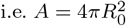, quantified the deviation from the spherical shape, with *v* = 1 for spheres. Shown are simulations with different T1 rate per cell *ν*. **e**, Cross-like initial in-silico organoid geometry relaxes into spherical shell via fluidization. Color-coded is the cell radius from the center of mass. **f,g**, The relaxation behavior toward a spherical shell upon fluidization for different T1 rates per cell *ν* (number of cells *N* = 450) and for different number of cells in shell *N* (for *ν* = 1.0). **h**, For a spherocylinder the reduced volume decreases with larger apical tension Γ_a_ (and larger apico-basal asymmetry), as the cylinder elongates (insets) (*N* = 277 cells, *ν* = 0.6). **i**, For an intestinal organoid the topology is inferred from the reconstruction. From this topology a mesh for the VM is created, for which the energy is then minimized. **j**, For reconstructed imaged intestinal organoids the simulated relaxation upon fluidization (*ν* = 2.0) is shown. Different organoids with different complexities are used, with exemplary initial configurations and configurations at *t* = 1000 (symbols) shown. Relaxation curves (f,g,h,j) are averages over multiple random simulations.

We verified this mechanism on cross-like simulated organoids (Fig. 4e) and measured the simulated relaxation time upon fluidization to find that shells relax faster with increasing T1-rate (Fig. 4f). We also find that increasing size indeed slows down relaxation (Fig. 4g). We further found that for slim spherocylindrical shapes, fluidization with apico-basal tension differences can even lead to elongation and further stabilization of preliminary buds (Fig. 4h). Including a luminal pressure leads to qualitatively similar behavior, albeit less pronounced (Extended Data Fig. E5).

To investigate this mechanism in an experimental context, we leveraged our organoid reconstruction to infer both tissue topology—who is neighbor to whom—and cell volumes, to import experimental organoid meshes into the VM (Fig. 4i, Supplementary Fig. S11). We find that both tissue topology and cell volumes already encode the organoid shape without apical tension patterning, if we also include the globally increased apical tension (Fig. 4i). No apical tension patterning is necessary to obtain the experimentally observed shape, consistent with our force inference result. Simulating organoid relaxation upon fluidization, we find that more complex organoids have a longer relaxation time (Fig. 4j, Supplementary Video 7). In fact recent tracking over 60 h in intestinal organoid buds has revealed that indeed the cells are dividing and rearranging, but no large-scale flows were found which could prevent the division-based structure formation [62]. We conclude that organoid shape stabilization is facilitated by the geometry, encoded by cellular volumes and the local topology of the polygonal tissue structure. With increasing size and complexity, a geometrical ratchet stabilizes organoid shape against fluidization.

### Line tensions develop over time and increase polygonal structure

In addition to calculating surface tensions and pressures, our approach also allows us to reconstruct line tensions, which act along the tangential directions of tri-interfacial junctions. The additional forces have to balance at vertices according to

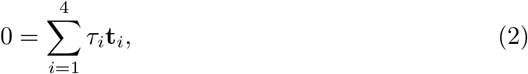

with force *τ*, which we will show to correspond to a line tension if present, and tangential vector **t**. Again, we normalize the relative tensions to yield an average of one. Typical values for the line tension should be around 10 nN [63]. Performing regularized inference on the corresponding line tensions, we obtain three-dimensional reconstructions (Fig. 5a) with broad distributions (Fig. 5b). We can conclude that the lateral line tensions are smaller than the basal and apical ones (Fig. 5c, Supplementary Video 8). At the same time, the apical junctions are much shorter (Fig. 5d), as to be expected from the round organoid shape and large thickness of the shell.

**Fig. 5.**
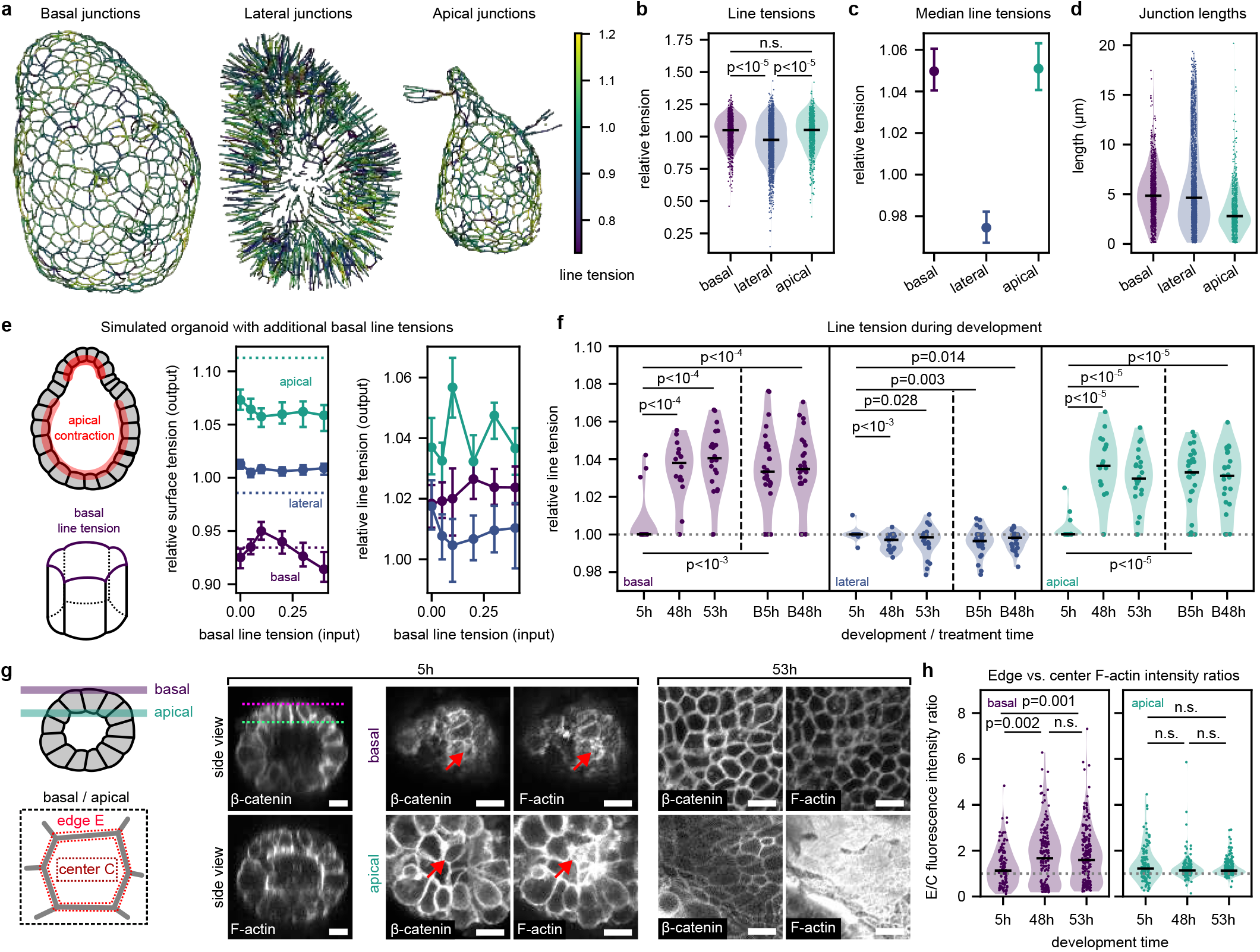
Force inference shows basal line tension increase concomitant with actin relocalization. **a**, Reconstructed basal, lateral and apical tri-interfacial junctions color-coded with inferred line tensions. Color-scale boundaries set to 5% and 95% quantiles. **b**, Distribution of inferred line tension by junction type. **c**, Medians of line tension distributions for different junction types with corresponding bootstrapped 95% confidence intervals. **d**, Distributions of the junction lengths by type. **e**, Simulated budded bubbly vertex model organoid with apical contraction (higher surface tensions) with additional basal line tensions (input). Results for inferred median surface tensions and line tensions (with 95% confidence intervals) for simulated organoid (output) with different assumed basal line tensions. Dotted horizontal lines are the ground truth medians. **f**, Basal, lateral and apical line tension median distributions for organoids at different developmental time points with and without blebbistatin treatment, conditions as in Fig. 3**i. g**, Imaging of basal and apical actin structures throughout development time. The membrane (*β*-catenin) marks the cell contours, with different localizations at cell center and edge of F-actin at the basal and apical sides. Arrows indicate exemplary cells with in-plane interfaces. Scale bars are 10 *µ*m. **h**, Quantification of basal and apical edge vs. center F-actin fluorescence intensity for cells that lie perpendicular to the imaging plane. As statistical test a Mann-Whitney-U-test was used. Black lines in violin plots indicate medians.

The finding that basal and apical line tensions are similar despite different junction lengths agrees with the notion that acto-myosin generated forces do not behave as elastic springs, but adjust their length by sliding filaments [13]. In detail, if the junction behaved like a spring, the force would increase with length, while for a pure line tension, the force would be independent of length. A plot of line tensions versus junction lengths confirms this relation and support the notion that the line tensions are generated by actomyosin forces, as are the surface tensions (Extended Data Fig. E6).

Reconstructing both surface and line tensions at the same time is challenging because mechanical force balance can be implemented both with and without line tensions. In particular, the experimentally found existence of a stronger apical surface tension makes it difficult to reconstruct the line tensions. To better understand the interplay between surface and line tensions, we performed simulations of a budded structured organoid in the BVM (Fig. 5e). Without basal line tensions in the simulations, we see that reconstructed apical line tensions are artificially increased. Basal and lateral line tensions are not separated, which is inconsistent with the observed increased basal line tension in the experimental data (Fig. 5c). Introducing a basal line tension into the simulations leads to a separation of the median basal and lateral line tensions (Fig. 5e), matching the experimental situation (Fig. 5c). In addition, a comparison of morphometric parameters such as cellular aspect ratios (see methods) is also consistent with the existence of additional basal line tensions (Extended Data Fig. E7).

Having established the importance and validity of the basal line tensions, we reconstructed them from the experimental data for the different stages of development (Fig. 5f). We find that basal and apical line tensions increase upon growth and symmetry breaking, consistent with no line tensions at 5 h (see methods and Supplementary Figure S3). While the apical line tension might reflect the same physical processes as the apical surface tension, the appearance of the basal line tension must reflect an increase in forces at the basal side that is not caused by cortical contractility.

Based on recent findings of special basal actin structures existing in fully formed intestinal epithelia [11, 64], we hypothesized that such structures could explain the increase in line tensions. Investigating F-actin localization over time, we found a strong interfacial signal basally at 5 h post-splitting, which becomes more junctional and polygonal-like over time (Fig. 5g). On the apical side, F-actin is organized in a more interfacial fashion throughout the entire time. Quantifying the edge versus center intensities, we indeed see a significant relocalization of basal F-actin in the medians of organoid samples (Fig. 5h; ratio increase from 1.1 to 1.6 from 5 h to 53 h), but also find a large variability between samples similarly to the observed line tension variability. On the apical side no statistically significant relocalization was found. Our results show that basal line tensions are effected by junctional actomyosin. They might be required to limit the opening angle of single cells, which otherwise might extrude [48, 65].

In conclusion, our new reconstruction pipeline for cell geometries and forces in organoids dominated by tensions now opens the door to disentangle the mechanical contributions of different force-generating components in epithelial systems, including engineered tissue models and colorectal cancer organoids. Such insight is crucial to unravel the physical basis of force generation and tissue-scale geometry in complex multicellular structures, both in physiological and disease conditions. The work presented here demonstrates that such an analysis is possible when combining experimental data with the appropriate physics concepts.

## Methods

### Organoid culture and preparation

Mouse small intestinal organoids were generated from the proximal part of the intestine of C57BL/6JRj mice. Animals were sacrificed for organ removal and used for scientific purposes in accordance with Section 4 TierSchG; and according to institutional licenses (DKFZ223-22) in compliance with German federal and state regulations. Mouse small intestinal organoids were generated as described previously [66]. Mouse small intestinal organoids were grown as described by Sato et al. [5], in mouse intesticult medium (Stemcell Technologies, #06005) supplemented with 100 mg/ml primocin (Invivogen, ant-pm-2). For experiments, mouse small intestinal organoids were seeded at a density of 20 crypts per 10 *µ*l BME type R1 (Biotechne, 3433-010-R1). For blebbistatin perturbation experiments, organoids were treated for the indicated amount of time to 10 *µ*M blebbistatin (Merck Milipore, 203389). At the end of the experiment, organoids were incubated in Cell Recovery solution (Corning, 354270) for 1 hour on ice to remove the matrix. For organoids exposed to blebbistatin, blebbistatin was included in the cell recovery solution to ensure maintained inhibition of actin-myosin contraction. Organoids were subsequently fixed in 4% formaldehyde for 30 minutes at room temperature.

For accurate staining of the membranes we first permeabilized the 0.5% Triton-X100 in PBS (Sigma-Aldrich, T9284) for 10 minutes at room temperature. Subsequently unspecific antibody binding was avoided by blocking the organoids for 1 hour at room temperature with blocking buffer consisting of 4%BSA, 0.1% Triton-X100, 0.1% Tween-20. Organoids were incubated overnight at 4°C in primary antibodies (Mouse-anti-Beta-Catenin, BD Biosciences, 610154, 1:500). Organoids were washed in 0.1% Tween-20 in PBS for 3 times before secondary antibody incubation for 1 hour at room temperature (Goat-anti-Mouse-AF488, Life Technologies GmbH, A11001, 1:500) and co-stained for F-actin (Phalloidin-TRITC, Sigma, P1951, 50 ng/ml) and DNA (Hoechst, Life Technologies GmbH, 62249, 1 *µ*g/ml). Organoids were washed in 0.1% Tween-20 in PBS for 3 times before a last wash in PBS before resuspension of the organoids in mounting buffer (2.5 M Fructose in 60% Glycerol) [67]. Organoids were mounted in mounting buffer in CellView chambers (Greiner, 543079).

### Confocal microscopy imaging and image analysis

Images were acquired on a Nikon AX laser scanning confocal microscope with a 25x Plan Apo Lambda silicone immersion objective (NA 1.05) with a voxel size of 0.25 × 0.25 ×0.5 *µ*m^**3**^. Raw data was deconvolved using the automatic 3D deconvolution software of the NIS Elements software, Version 5.40.20. For the quantification of F-actin intensities the raw imaging data was used. Cells were identified which lie perpendicular to both the basal and apical imaging planes and edge outlines were drawn based on the *β*-catenin membrane signal using the ‘freehand’ line tool in ImagJ with line width 3. Then z-stacks were acquired with six slices, giving a thickness of 3 *µ*m. For each organoid the matching basal, middle and apical sections from 3 cells were analysed. Images in Fig. 5g within the respective organoid display same relative intensity per imaged structure, with the exception of apical *β*-catenin, where contrast was linearly increased to increase visibility of edges. Organoids without perfectly aligned cells or with aligned cells undergoing cell divisions were excluded from the analysis.

### Bubbly vertex model simulation of in silico organoids

In the classical vertex model individual cells are described by polyhedra with apical, basal and lateral polygonal interfaces, subject to surface tensions. The interfaces are described by vertex nodes which define the (not-necessarily planar) polygons. We assume the same network topology on the apical and basal side, i.e. the lateral interfaces are quadrilateral. The energy reads

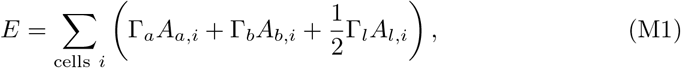

with apical, basal and lateral surface tenseions, Γ_*j*_ for *j* = a, b, l, respectively. The (summed) interfacial areas *A*_*j*_ for these interfaces then contribute to the energy, where we account for lateral membrane double counting via a factor of 1*/*2. Following Drozdowski and Schwarz [48] and using OrganoidChaste [68], we minimize this energy for a VM configuration via overdamped dynamics for the vertex nodes defining the polyhedra with simultaneous simulated annealing in the rate of cell neighbor exchanges (T1 transitions). For the overdamped relaxation we consider a force on vertex node **r**_*k*_

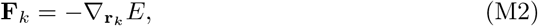

and consider the overdamped equation of motion

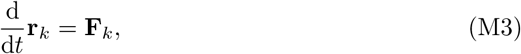

which is integrated using Euler stepping. The configuration is then exported to SurfaceEvolver [50] and subsequently minimized to obtain the Bubbly Vertex Model. In SurfaceEvolver luminal volume constraints were used in simulated in silico organoids when performing force inference in order to unfold collapsed/small interfaces to obtain interface sizes comparable to experiments (Supplementary Figure S4). In particular, in Fig. 2d we chose *V*_lumen_*/V*_VM_ = 3 for the bubbly lumen volume compared to the (not bubbly) vertex model lumen volume; in Fig. 3h *V*_lumen_*/V*_VM_ = 1; in Fig. 5i *V*_lumen_*/V*_VM_ = 2; and in extended data Fig. E7b,c *V*_lumen_*/V*_VM_ = 1. For the simulated organoids in Fig. 5i the distributions from which the surface tensions are drawn are Gaussian with a standard deviation of 0.005 and means of 1 for lateral; 1.0 (neck) 1.15, and 1.35 (buds) for apical; and 1.0 (neck), and 0.95 for basal interfaces.

For the analysis and force inference in in silico organoids we used the software Paraview [69]. The resulting in silico organoids were sliced to create a 3D image stack, which was segmented using an appropriate Ilastik model [46] and reconstructed with *ForceIn3D* [70]. The voxels in which the two-dimensional interfaces lie were fully colored, effectively yielding a sharp box-shaped point spread function.

In unstructured spherical in silico organoids, we found line tensions to be over-regularized. This yields a strong peak at one in the BVM, similarly to what was observed in the 5 h unstructured case (Supplementary Figure S3). Very small inferred deviations of line tensions around one therefore correspond to unstructured spherical organoids without real line tensions.

### Image segmentation pipeline

The microscopy image stacks of organoids contain three channels: magenta (F-actin), green (*β*-catenin) and blue (nucleus). They are preprocessed by performing depth-correction adjusting the intensity loss for deeper imaging planes, gamma correction and image registration if necessary. In addition, images are interleaved to obtain cubic voxels with an edge length of 250 nm. Figure 1d shows the segmentation pipeline: the green and red channels are separated (1) and an auxiliary segmentation is created with Cellpose [44, 45] (2), using a model trained with additional experimental organoid data. Through edge-filtering an edge probability map is calculated (3). Together with the full image (including the blue nucleus channel) these are segmented using ilastik [46] (4), yielding the final three-dimensional segmentation. For nucleus segmentation depth and gamma corrections were performed on nuclear imaging data with subsequent segmentation via Cellpose [44, 45] and ilastik [46]. Details are given in Supplementary Note 1.

Our procedure yields satisfying segmentation results. Nonetheless, segmentations are not perfect. We found that in a few cases apical or basal interfaces are missed by our segmentation pipeline, which leads to the connection of both a lateral, and a basal or apical interface into one point cloud. To correct for this, and assuming that apical and basal interfaces do not connect the lumen with the outside (only lateral interfaces do), these interfaces were identified as segmentation errors and excluded from the statistical analysis. For some interfaces stemming from segmentation errors such automatic removal is not possible (cf. Fig. 3a and Fig. 5a). Since these spurious interfaces and junctions generally mix lateral data into apical or basal data, this does not influence our statistics. Such wrong classifications lead to a lowering of reported statistical effects as deviations from the average values are considered.

### Three-dimensional non-parametric reconstruction of junctions, interfaces and organoid morphology

We adopt a point cloud based methodology as first suggested by Xu et al. [34], which we modified to yield reliable results for our data. For the tri-interfacial junction reconstruction the voxels with three neighboring cells in a 19-point stencil environment (next-nearest neighbors, Supplementary Fig. S5) are compiled into a voxelated representation. For this then the algorithm by Lee [71] was used, with the modification that an initial estimate of the tangential space around each point is done via principal component analysis (PCA). Then a neighborhood size is determined on which the curve is mostly linear and in that neighborhood a quadratic function is fit in the osculating plane of the curve (Supplementary Fig. S6). The points are then projected onto that locally fit curve. For details see Supplementary Note 1.

Similarly, we determine the interface voxels via a 19 point stencil by determining voxels with two neighboring cells and additionally including junctional voxels at the boundaries. For the reconstruction of the interfaces we estimate the normal vector via PCA, fit a quadratic surface on a neighborhood where the interface can be estimated to be in a planar regime (generalizing the algorithm for one-dimensional curves), and project the point on the locally reconstructed interface (surface thinning). The normals are then corrected to match the reconstructed surface. Curvatures are calculated on the thinned interface (Supplementary Note 1).

Organoids can also be considered as a surface with finite thickness, like the cellular interfaces. Thus we use the interface reconstruction on voxels belonging to cells, to determine the mid-plane and its curvatures. The surface thinning algorithm can be used iteratively to yield better thinning results, which we did in the morphological reconstruction (Supplementary Note 1). Note that the algorithm can only determine the mid-plane well in the case of radii of curvatures larger than the thickness, which is the case in the reconstructed simulated and experimental organoids in this study.

All the aforementioned reconstruction methods were verified on corresponding in silico test cases (Supplementary Note 1 and Supplementary Figs. S7-S9).

### Force and pressure inference

Force balance from surface tensions, Eq. (1), and from line tensions, Eq. (2), and the Young-Laplace law for pressure differences are linear equations for **x** – the vector of surface tensions, line tensions or (rescaled) pressures, respectively. The resulting matrix equation reads

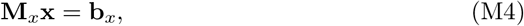

with right hand side of the equations **b**_*x*_. To solve for the unknown **x** we use a least squared error (LSE) approach, from which we obtain, using the speudoinverse,

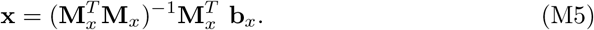

Since the pseudoinverse can be ill-conditioned due to the noisy ill-posed nature of the inverse problem and due to segmentation errors, we perform Thikonov regularization. The LSE minimizer, which serves as an estimator for the quantity to infer, **x**, then reads

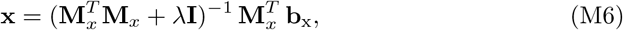

with identity matrix **I**. The regularization parameter λ is chosen as such that it minimizes Allen’s predicted residual error sum of squares (PRESS) statistic in Leave-One-Out cross validation (LOOCV) [51]. For details see Supplementary Note 2. Inference robustness against variations in the regularization parameter *λ* and against incomplete reconstruction information from interface removal was validated on in-silico data (Supplementary Fig. S10).

### Dynamical simulation and geometry extraction of organoids

To simulate organoid dynamics in the limit of flat interfaces (the classical VM) we use *OrganoidChaste* [48]. Assuming constant apical, basal and lateral surface tensions *ϒ*_a_, *ϒ*_b_, and *ϒ*_l_, and line tensions, *τ*_a_, *τ*_b_, *τ*_l_, respectively,the Hamiltonian reads

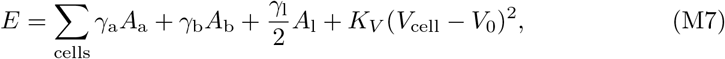

with (cell-specific) apical, basal and lateral total areas, *A*_a_, *A*_b_, and *A*_l_, and edge lengths *𝓁*_a_, *𝓁*_b_, *𝓁*_l_, respectively, elastic volume modulus *K*_*V*_, cellular volume *V*_cell_ and (cell-specific) target volume *V*_0_. For simulations we assume *K*_*V*_ = 20.0 to effectively have a stiff volume conservation constraint. Vertices defining the geometry are moved in an overdamped fashion according to the gradient of this energy. Active cell rearrangements are implemented through random selection of lateral interfaces, which are collapsed to a line and resolved orthogonally to perform a T1 transition. To include a pressure, we consider d*E* = *p*dV and compute the force from this by considering luminal volume changes. For growing organoids, stem cells divide into one stem and one transit amplifying (TA) cell (only one generation of TA cells). A TA cell divides into two differentiated cells. We assume that cells double their volume for *t*_sim_ = 25 prior to division and stay at *V* = 1 for *t*_sim_ = 25 after division, i.e. division occurs every *t*_sim_ = 50.

To extract the geometry of a reconstructed experimental organoid a voxelated organoid mask is created. This mask is smoothed (Gaussian filter 10 voxel standard deviation), downsampled (factor 4) and then using marching cubes [72] a 3D triangular mesh of the basal (outer) organoid surface is created. Triangles are assigned to basal interfaces via minimum distance. Then a dual mesh on the surface is created, yielding a Voronoi-like coverage of the surface. Resulting polygonal interfaces stemming from the same basal interface are merged and mesh artifacts like inclusions and interfaces without at least three neighbors are removed through merging into neighboring polygons. The resulting mesh is simplified to only save corner nodes which separate three interfaces (Supplementary Figure S11). This topology is then read into *OrganoidChaste*, where the monolayer mesh is created with a small thickness, which is relaxed into the real monolayer thickness. Space is non-dimensionalized by the median cellular volumes and we found a median dimensionless height of *h* ≈1.9 in all organoids used for simulation. Cellular volumes are set by taking the polygonal area times the uniform height. For organoid relaxation upon fluidization (Fig. 4j) only topology is considered (cell volumes are spatially homogeneous, no apico-basal tension asymmetry). Mesh inference code was written with help from large language model Claude and verified and tested manually.

### Mouse donor normalization of inferred tensions

We performed experiments from organoids which were created from initial samples from 4 different mouse donors. We observed deviations in the global median tensions between different mouse donors (Supplementary Figure S2). Due to the large number of lateral interfaces and the averaging of tensions to 1, the median lateral tensions are very close to 1. To account for the mouse-to-mouse variability, we perform a donor-specific normalization, rescaling the average apical deviation from 1 (the lateral tensions) to 1. We thus normalize tension via

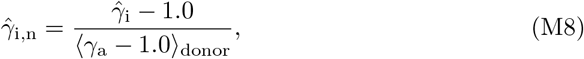

where we consider deviations from one, rescaled by the average deviation of the apical tension (*ϒ*_a_) averaged over the mouse donor,⟨ . ⟩_donor_. Here 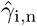 is the normalized tensions of interface *i*, which can be either apical, basal or lateral. For this rescaling we only consider the 53 hour control experiments when also different additional conditions are considered.

### Comparisons of cell heights and aspect ratios in the vertex model

To quantify how well tissue shape can be understood based on surface tensions, i.e. the vertex model and the bubbly vertex model, we compared experimental aspect ratios of cells, with the corresponding theoretical aspect ratios we would expect from a hexagonal vertex model (extended data, Fig. E7).

The experimental aspect ratio of a cell in the segmented data can be defined through the cell height *h* and the mid plane area *A*_mid_ to be

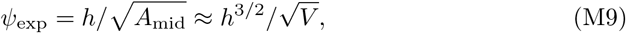

where we use approximation for the volume *V* = *A*_mid_*h*. Cell heights are determined from the segmentation by calculating the average position of the apical and basal interfacial voxels in space and considering the average points’ distance.

To compare to a theoretical estimate, we consider a polyhedral representation of single cells in the vertex model, described by Eq. (M1). The cell height in the case of hexagonal cells (the close packing solution of the plane) can be determined by minimizing the energy as a function of height [73] to be

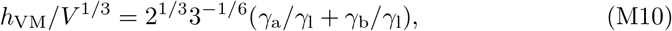

with apical, basal and lateral (normalized) surface tensions *ϒ*_a,b,l_, respectively. Considering lateral sides with an averaged lateral surface tension, i.e. we replace the lateral tension by the average of the *n* lateral tensions, thus yields as theoretical expectation for the aspect ratio

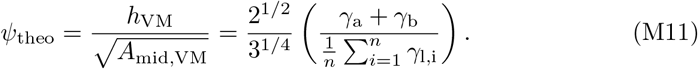

Note that this aspect ratio is the theoretically expected ratio for hexagonal vertex model cells, i.e. we neglect interfacial curvatures. Our comparison to the bubbly vertex model simulations show, however, that the average aspect ratio in the bubbly vertex model matches this theoretical value.

Assessing the ratio of the two aspect ratios allows a quantification of the deviation from the surface-tension-based morphology. We simulated budded organoids with increasing basal line tensions (extended data, Fig. E7b,c) and found that in the BVM the aspect ratio increases when a basal line tension is introduced, yielding an aspect ratio deviation (i.e. the ratio of experimental and theoretical aspect ratios) greater than one.

To theoretically determine the height of a hexagonal cell with an additional basal line tension, we consider the modified energy of a single hexagonal cell in a flat monolayer

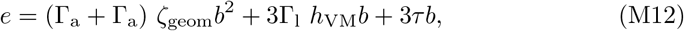

with basal edge length *b* and geometrical constant ζ_geom_ = 3^3*/*2^*/*2. Denoting the height without line tension as *h*_0_, the corresponding height *h* which minimizes this energy for a given *τ* is given by

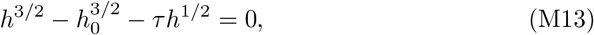

and the corresponding aspect ratio deviation then reads 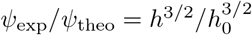. The cell height and the aspect ratio deviation increase when *τ* is increased, explaining the corresponding simulated results.

Note that luminal volume constraints, in analogy to luminal pressures, lead to stretching and therefore a lowering of the aspect ratio (Supplementary Figure S4).

### Nuclear shape calculation

For each nucleus a point cloud from the segmented voxels is created. The long axes are determined using principal component analysis (PCA) on the resulting point cloud. The maximum distances in the point cloud in directions of the PCA are used to estimate the orthogonal lengths *v*_1_ *≥ v*_2_ *≥ v*_3_.

### Statistics and reproducibility

All experiments were performed with samples obtained from at least 4 mouse donors, where from two mice at least two full technical replicates were collected, i.e. the experiment was performed independently at least two times for these mice with all conditions in the dataset (Supplementary Table S1). The data from multiple independent experiments was pooled in the analysis. The total number of necessary samples per experimental group was estimated from similar previous experiments [6]. For the developmental time course (Fig. 3j,k; Fig. E4d,e; Fig. 5k; extended data Fig. E7e,f,g) the number of organoid samples are *n*(53 h) = 22, *n*(48 h) = 20, *n*(5 h) = 20, *n*(B5 h) = 28, *n*(B48 h) = 25. For the F-actin intensities (Fig. 5h) the number of basal interfaces are *n*(53 h) = 15, *n*(48 h) = 20, and *n*(5 h) = 25; and of apical interfaces are *n*(53 h) = 14, *n*(48 h) = 18, and *n*(5 h) = 24. Simulations of in-silico organoid shape relaxations (Fig. 4, extended data Fig. E5) were averaged over *n*_sim_ = 4 simulations. For the edge versus center F-actin intensity analysis (Fig. 5h) in total 59 organoids were analysed comprising 3 replicates and 3 biological donors with 3 sampled cells per organoid, leading to a total of *n*_cells_(5 h) = 135, *n*_cells_(48 h) = 171, and *n*_cells_(53 h) = 225 cells. All statistical tests were performed with the python package SciPy.

## Data availability

The trained Cellpose model, the trained ilastik models, the summary data of all analyzed organoids, the segmented exemplary organoid used in this study, with the corresponding force inference data, and the scripts to analyze and visualize the force inference data is available on heiDATA, Heidelberg’s open research data repository, at https://doi.org/10.11588/DATA/H6EW4G.

## Code availability

The force inference code, *ForceIn3D*, is available on Github [70]. The code for simulation of the bubbly vertex model, *OrganoidChaste*, with the corresponding simulation files, is also available on Github [68].

## Acknowledgements

O.M.D., M.B. and U.S.S. acknowledge support by the Max Planck School Matter to Life, with funding by the German Federal Ministry of Education and Research (BMBF), the Dieter Schwarz Foundation, and the Max Planck Society. M.B. and U.S.S. acknowledge support by the Deutsche Forschungsgemeinschaft (DFG, German Research Foundation) through the cluster of excellence 3DMM2O (EXC 2082/1-390761711 and EXC 2082/2-390761711). The authors acknowledge support by the state of Baden-Württemberg through bwHPC and the DFG through grant INST 35/1597-1 FUGG, and the data storage service SDS@hd supported by the Ministry of Science, Research and the Arts Baden-Württemberg (MWK) and the DFG through grant INST 35/1503-1 FUGG. U.E. acknowledges Nikon Europe for support of the Nikon Imaging Center at Heidelberg University, CellNetworks core technology platform and the CRC 1324. Research in the lab of M.B. was supported by the CRC 1324 (DFG). U.S.S. is member of the Interdisciplinary Center for Scientific Computing (IWR) at Heidelberg.

## Author contributions

O.M.D. and K.E.B. contributed equally to this work. In particular, O.M.D. developed the software and performed the simulations. O.M.D analyzed the data with contributions from K.E.B.. K.E.B. developed the experimental protocols and performed the experiments. K.E.B. imaged the organoids with help from U.E.. O.M.D. and K.E.B. designed the experiments together. O.M.D and U.S.S. developed the theoretical model. U.S.S. and M.B. supervised the work. O.M.D. and U.S.S. wrote the paper together with help from K.E.B. and with input from all other authors.

---

**Fig. E1.**
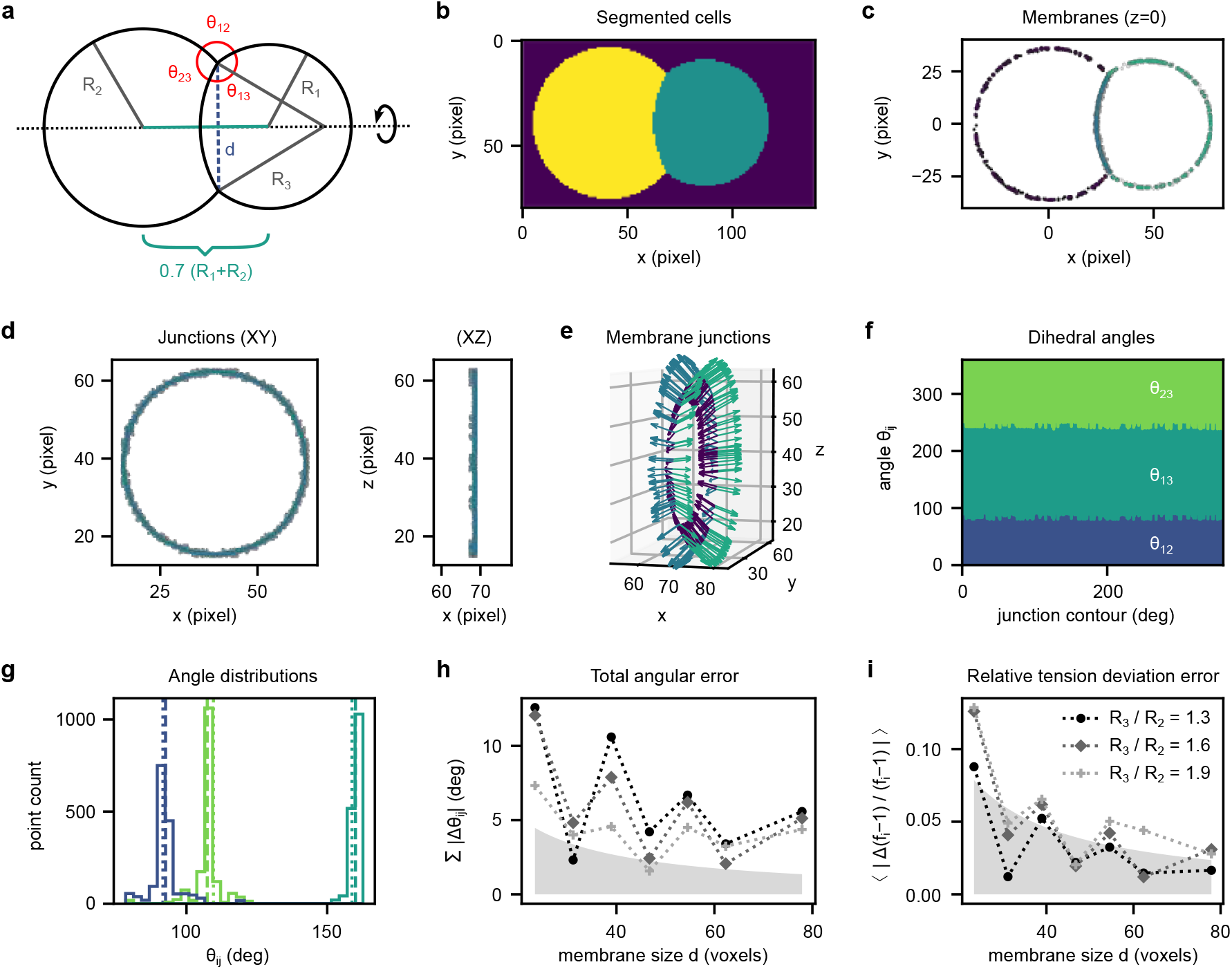
Force inference results for a prescribed in silico two-cell system. **a**, Schematic depiction of the in silico two-cell system used for testing the force inference method. Two spherical cells with radii *R*_1_ and *R*_2_ are separated by a spherical-cap interface with radius *R*_3_ and opening diameter *d*. The angles *θ*_*ij*_ allow for a reconstruction of relative tensions. **b**, A cut (*z* = 0) through the voxelated cell segmentation for the two-cell system. We consider a system with *R*_1_ = 30, *R*_2_ = 1.2·*R*_1_, *R*_3_ = 1.3 · *R*_2_ in **b-g. c**, Cut through the point cloud reconstruction of the interfaces with initial (black) and thinned (color) point clouds. **d**, The tri-interfacial junction reconstruction with initial (black) and thinned (color) point clouds, projected into the XY and XZ planes. **e**, Three-dimensional reconstruction of the junction with tangential vectors of the interfaces. **f**, Dihedral interface angles along the junction contour. **g**, Distribution of the dihedral angles, color-coded as in **f**. Dotted lines are the theoretically expected angles from the given configuration and dashed lines the means of the distribution. **h,i**, For different resolutions, corresponding to interface sizes *d*, the errors in force inference are determined. Radius *R*_1_ is varied with *R*_2_ = 1.2 · *R*_1_ and different *R*_3_*/R*_2_ in color. The total angular error **(h)** and the relative tension deviation errors **(i)** are shown, see Supplementary Note 3 for details. Shaded in grey is an estimate of the error range from the voxel resolution.

**Fig. E2.**
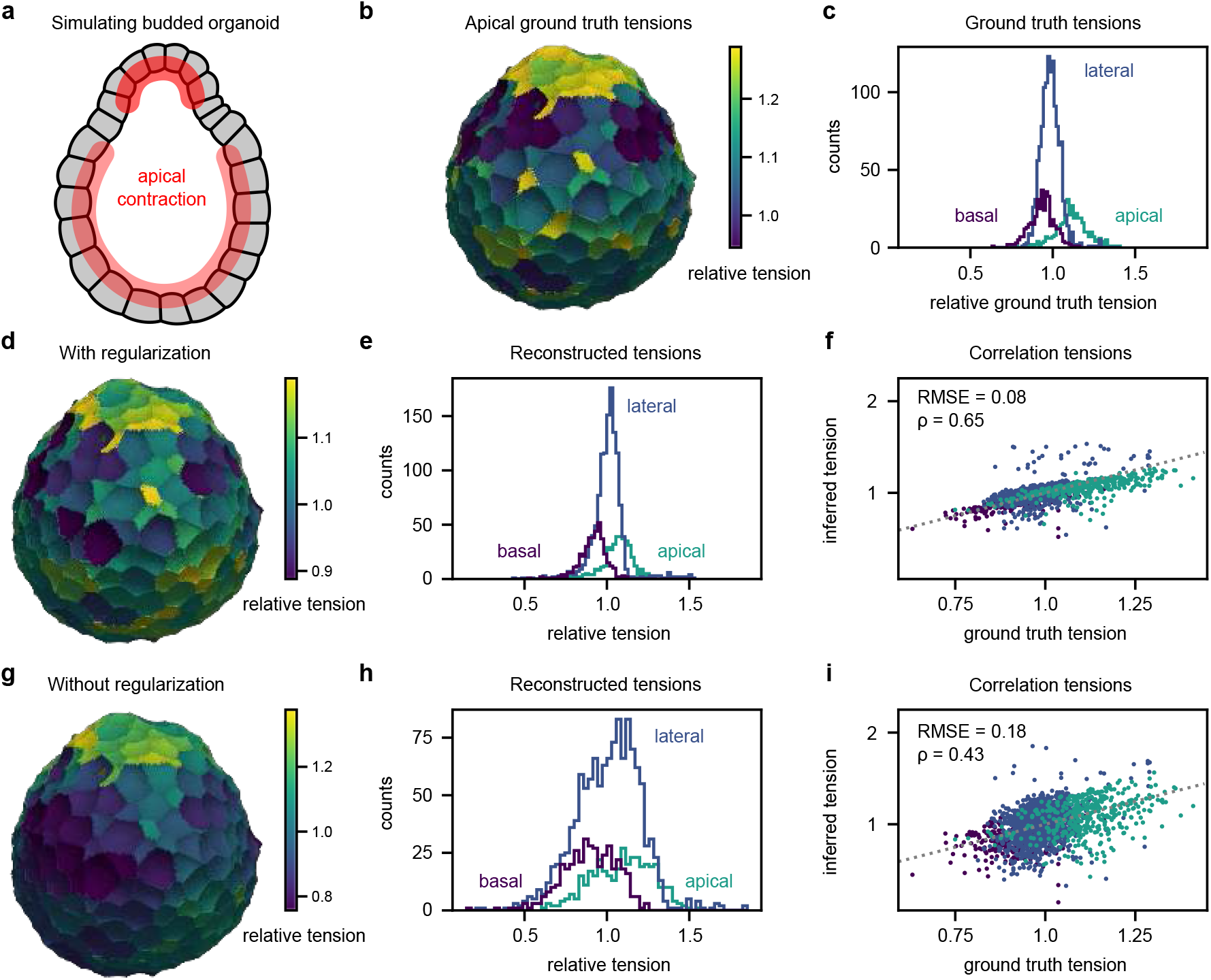
Force inference of a budded pressurized in silico organoid with and without regularization. **a**, Schematic depiction of apical contraction (red) effecting higher apical tensions in the buds than in the spherical rest, resulting in spontaneous curvature and leading to a budded shape. **b**, Simulated pressurized in silico organoid with the bubbly vertex model with 400 cells, luminal volume *V/V*_VM_ = 2.0 (see methods) and apical tension patterning leading to a budded shape, color coded with relative tensions. **c**, Real ground truth tensions in the simulated organoid. **d**, Reconstructed interfacial tensions with Thikonov regularization in the inverse force inference step. **e,f**, Distribution of reconstructed tensions with regularizations, and correlation with real ground truth tensions with root mean squared error (RMSE) and Pearson correlation coefficient *ρ*. **g**, Reconstructed interfacial tension without regularization. **h,i**, Distribution and correlation without regularization. All tensions color-coded with range between 5% and 95% quantiles. The distributions from which the real tensions are drawn are Gaussian with a standard deviation of 0.015 and means of 1 for lateral, 1.15 and 1.3 for apical, and 0.95 (buds) and 1.0 for basal faces.

**Fig. E3.**
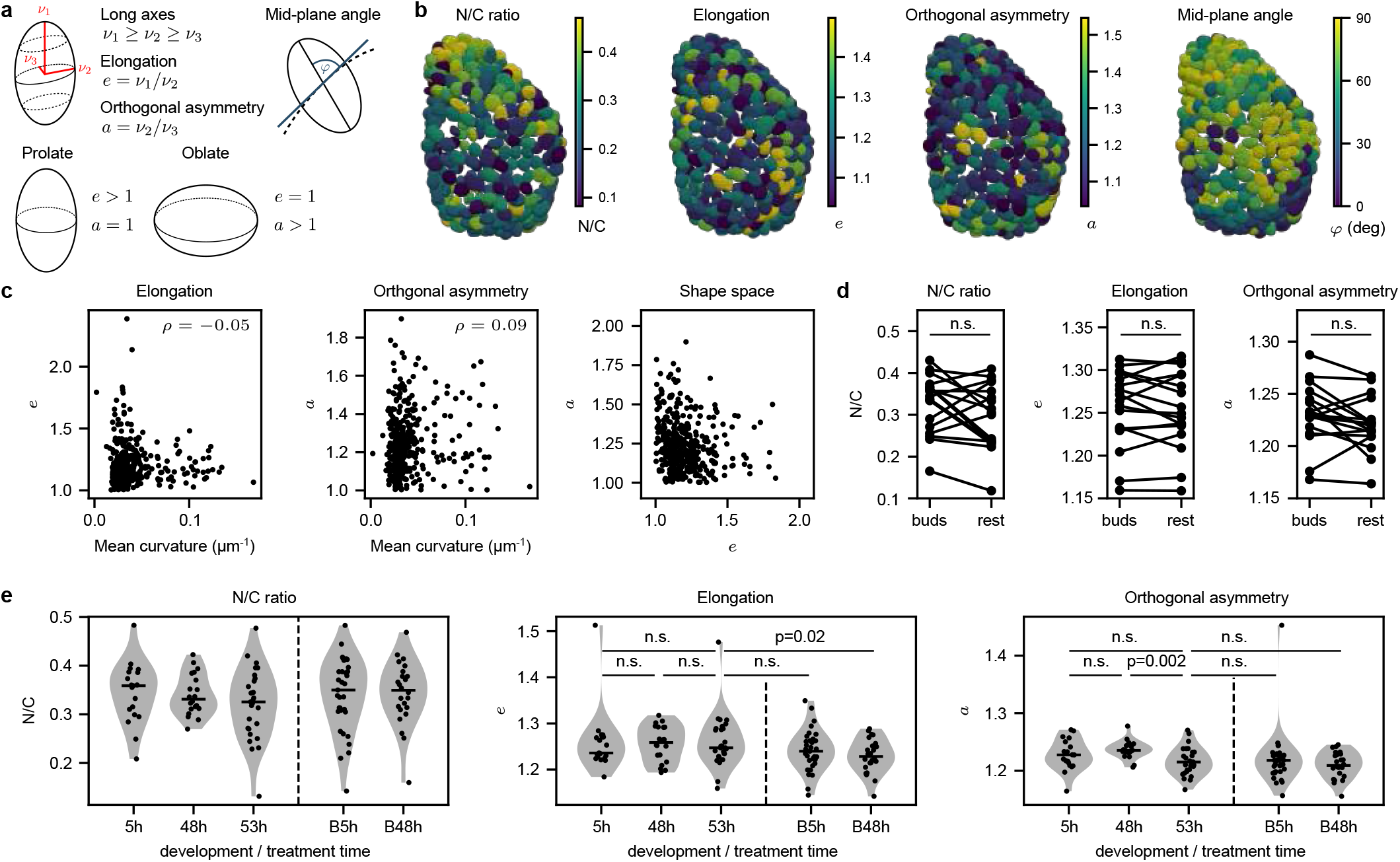
Nuclear shapes in budding organoids not strongly affected by morphological changes. **a**, Nuclear shapes are quantified by determining the long axes and calculating the elongation e, the orthogonal asymmetry a and the mid-plane angle φ. **b**, Nuclear-to-cytoplasm volume ratios (N/C), elongations, orthogonal asymmetries, and mid-plane angles for budded organoid color-coded with range between 5% and 95% quantiles. No clear patterning is visible except for N/C, which is slighly increased in the bud. **c**, Elongations and orthogonal asymmetries as functions of midplane mean curvature with correlation coefficient ρ. Nuclei are shown in (a, e) shape space. No clear dependence on curvature is visible and many nuclei have anisotropic shapes (a > 1, e > 1). **d**, Comparison of different nuclear shape quantifiers between budded and non-budded regions in 53 h organoids. No significant differences are found, but in some organoid N/C and orthogonal asymmetry are higher in buds. **e**, Comparison of different nuclear shape quantifiers across development time. No differences in time were found except for a slight drop in orthogonal asymmetry after 48 h, indicating a smaller anisotropy after budding. As statistical tests Wilcoxon (**d**) and Mann-Whitney-U-tests (**e**) were used. Black lines in violin plots indicate medians.

**Fig. E4.**
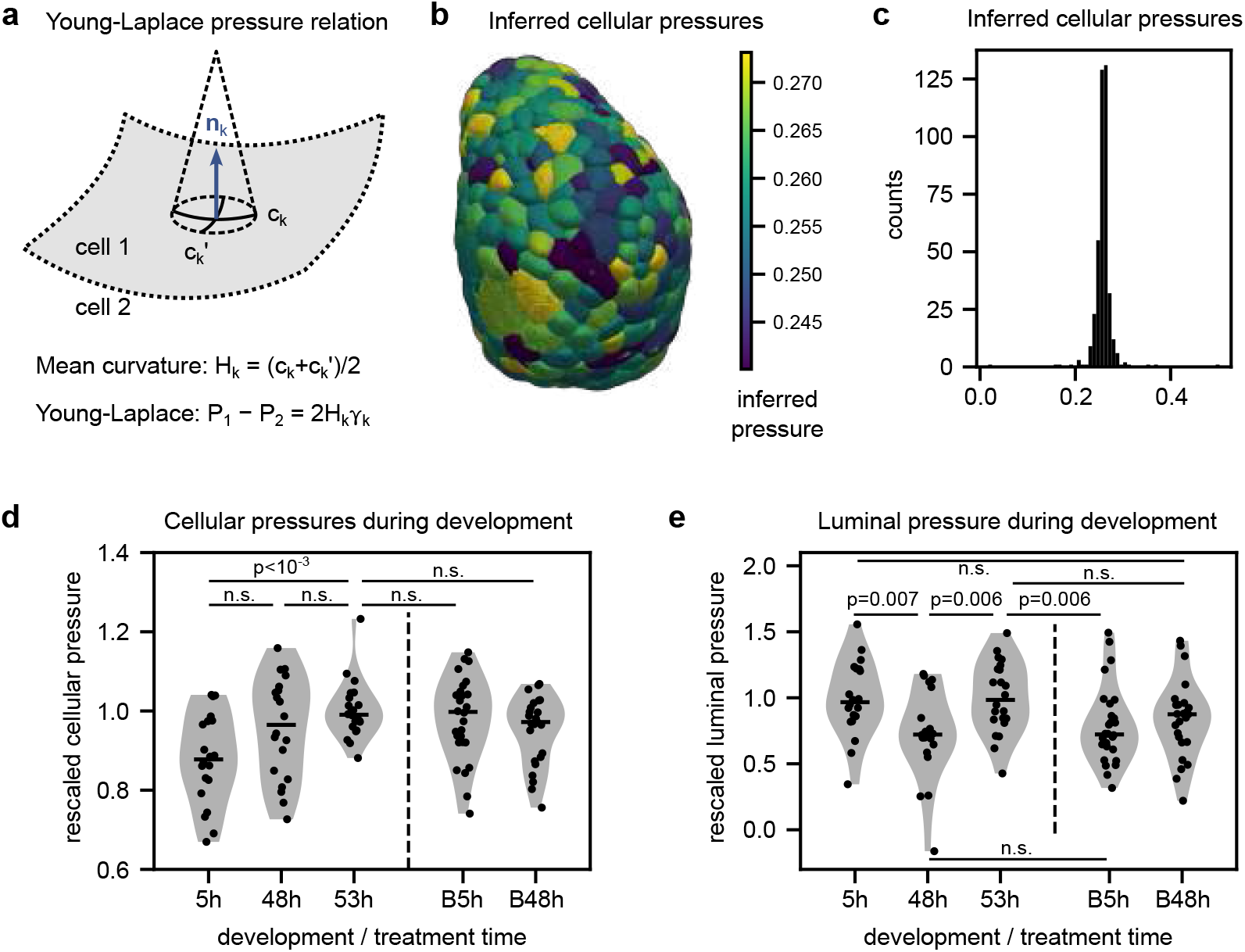
Cellular and luminal rescaled pressure inference. **a**, Across an interface k between cells 1 and 2 the mean curvature 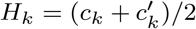, with principal curvatures 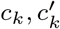, in the direction of the normal n_k_ and the interfacial tension γ_k_ are related to the difference in pressures P_1,2_ via the Young-Laplace law. Inference via known curvature and tension thus yields pressure differences rescaled by the (unknown) average surface tension. **b**, Rescaled cellular pressures for budded organoid colorcoded with range between 5% and 95% quantiles. **c**, Cell pressure distribution from organoid in **e. d,e**, Cell pressure median and luminal pressure distributions for organoids at different developmental time points with and without blebbistatin treatment. Conditions as in Fig. 3**i**. As statistical test a Mann-Whitney-U-test was used. Black lines in violin plots indicate medians.

**Fig. E5.**
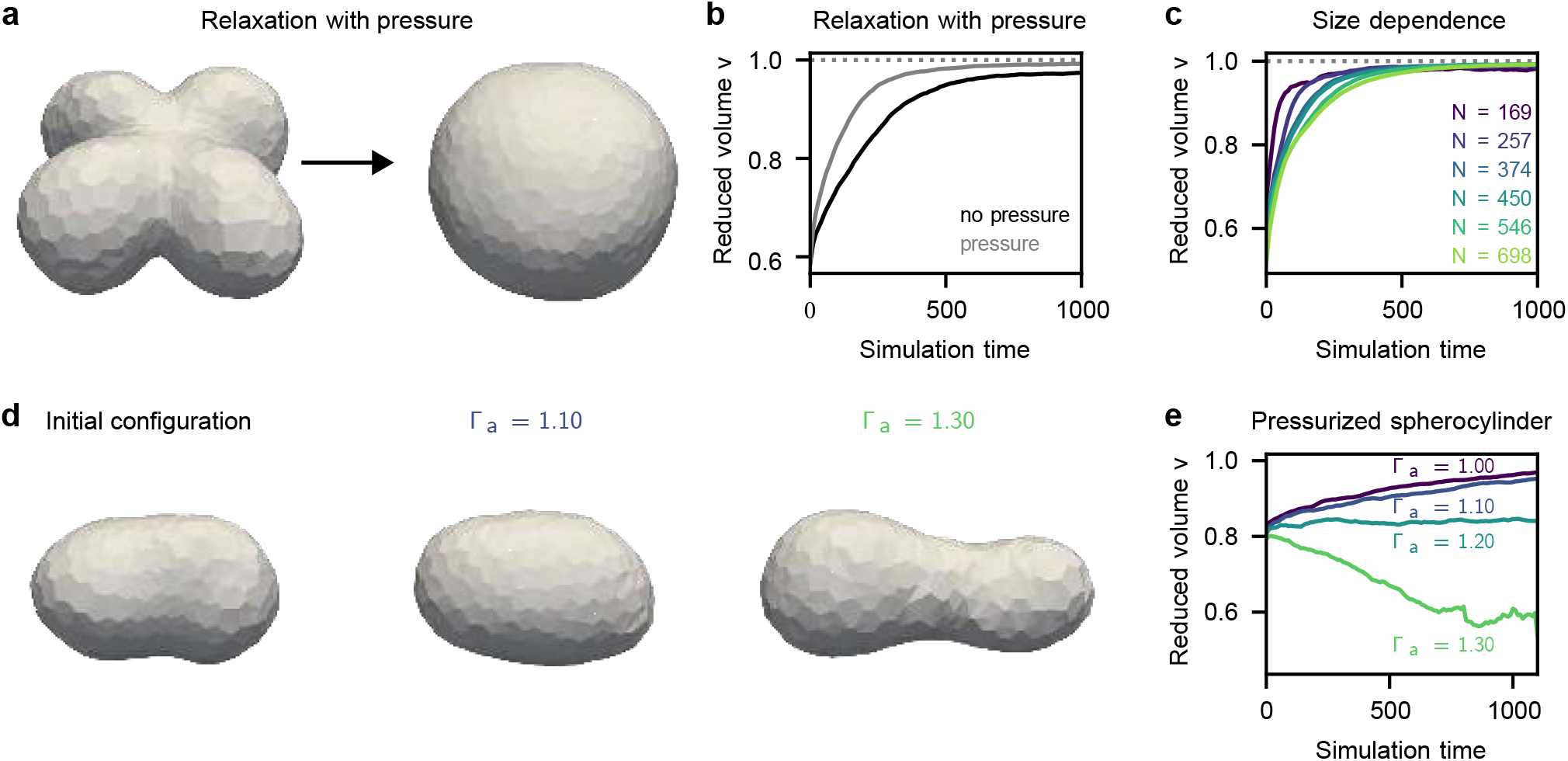
Geometrical ratchet behavior with a pressurized lumen. **a**, For cross-like initial in-silico geometries with luminal pressure the organoid relaxes upon fluidization (*N* = 277, *P* = 0.05, *v* = 1.0). **b**, Comparing relaxation without (black) and with pressure (grey), we see similar relaxation behavior. **c**, Similarly as in the unpressurized case shells relax slower when they are larger. However with pressure this effect is not as strong. **d**, Spherocylinder with luminal pressure (*P* = 0.05) relaxation starts from an elongated shape and relaxes to a more spherical configuration for small apico-basal tension difference, i.e. small Γ_a_ for Γ_b_ = Γ_l_ = 1, and to an elongated shape until the shape splits for large Γa. **e**, For different apical tensions Γ_a_ the reduced volume either increases or decreases, resulting from the elongation instability. Relaxation curves (b,c,e) are averages over multiple random simulations.

**Fig. E6.**
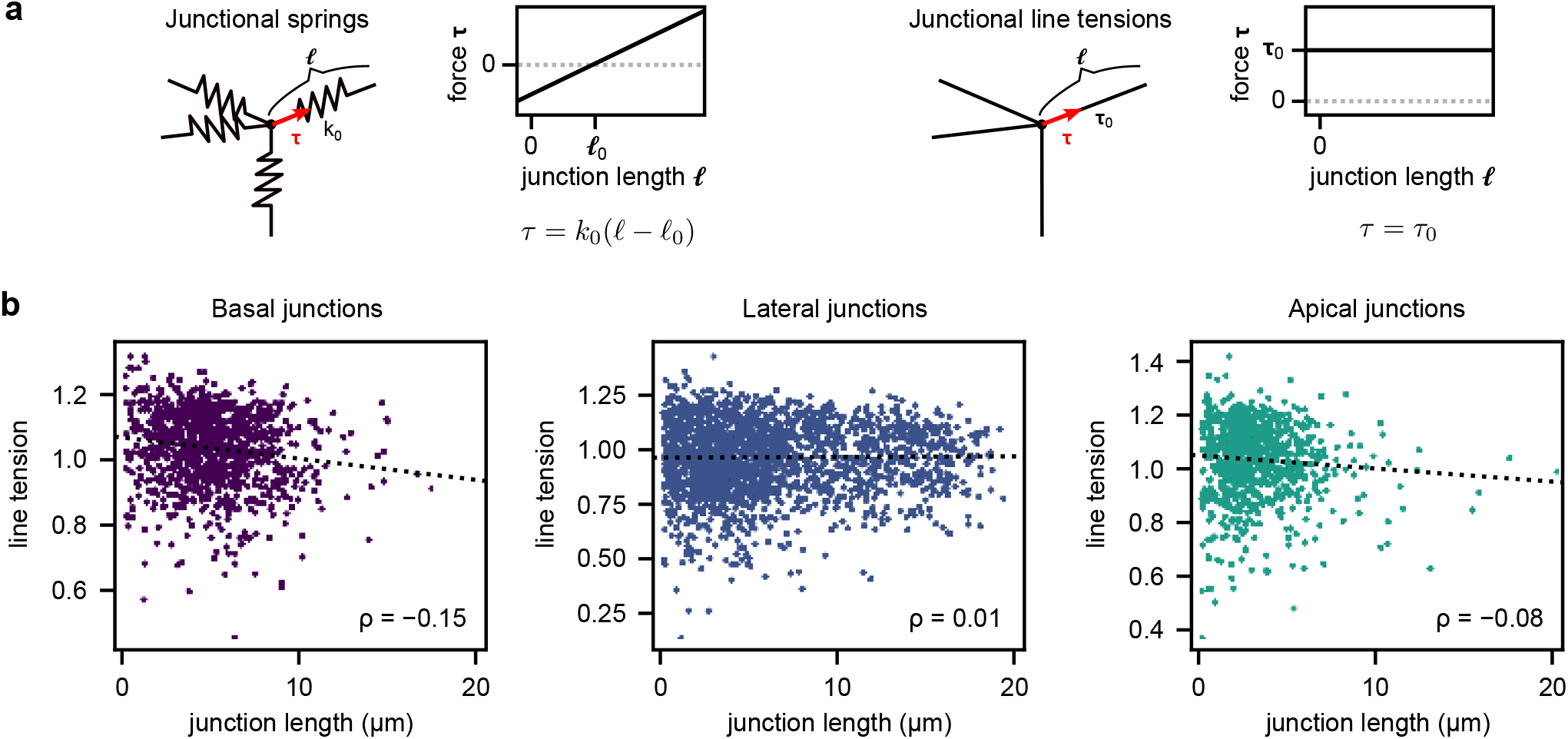
Comparison of spring-like and tension-like tri-interfacial junction models. **a**, Models for different origins of tangential junction force contributions with corresponding single-junction force extension curves: junctional springs, which have a linear curve with offset for an individual junction, and line tensions, which have a constant force. **b**, Inferred line tensions as function of the junction length with corresponding correlation coefficient *ρ* for organoid shown in Fig. 5a-d. The negatively correlated forces are consistent with line tensions in a multicellular structure, where the cells adopt shapes minimizing their total energy, leading to smaller junctional lengths for larger tensions.

**Fig. E7.**
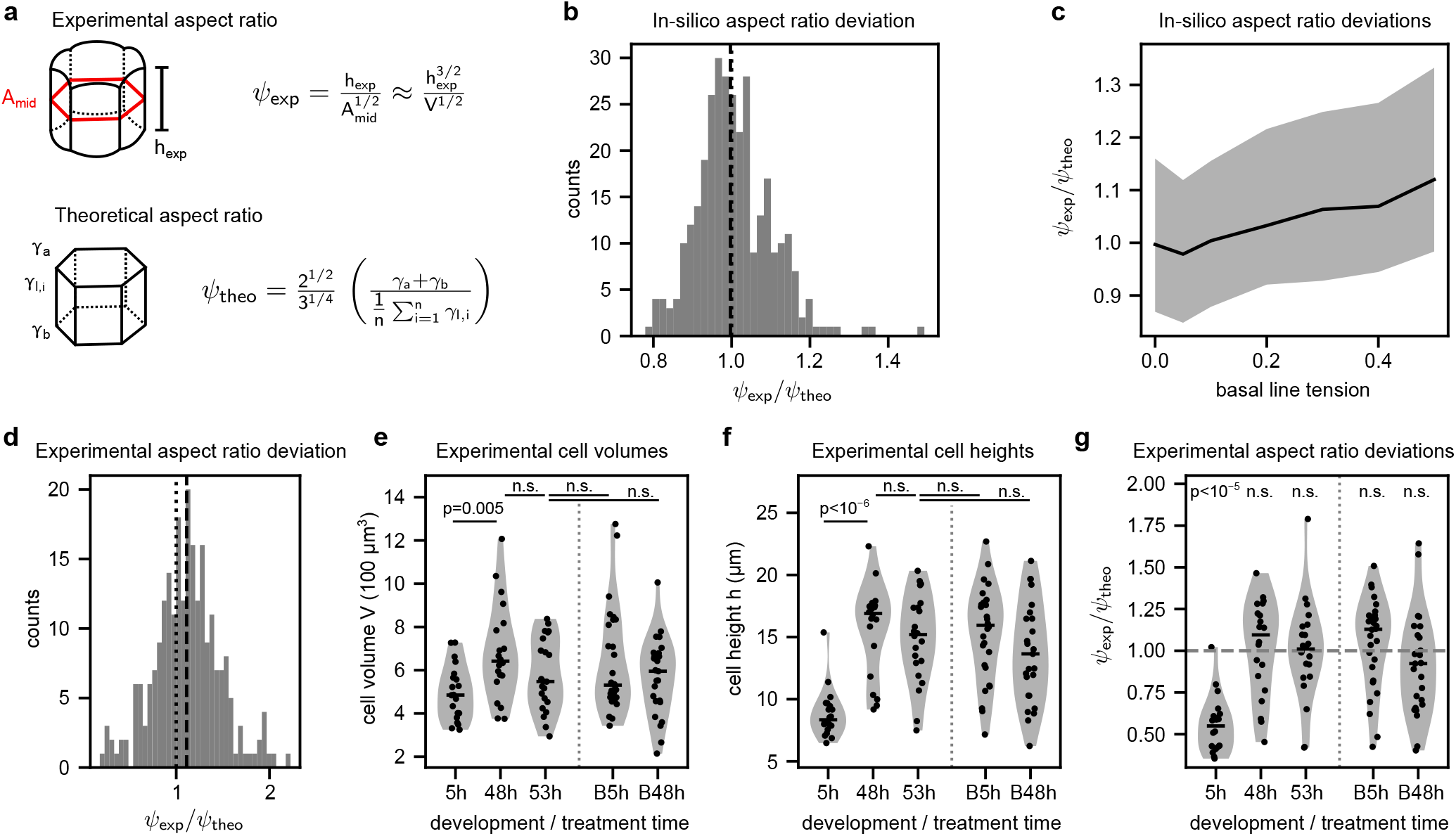
Morphometric comparison to the bubbly vertex model. **a**, Definition of the experimental aspect ratio, which can be computed approximately from the height and volume, and of the theoretical aspect ratio (see methods), which is obtained from the surface tensions *γ*. **c**, Distribution of aspect ratio deviations in simulated organoid (cf. Fig. 3**h**), which shows agreement between experimental and theoretical aspect ratios. **d**, Aspect ratio deviations for simulated organoids with an additional basal line tension. Full line is median with shaded region showing the 5% to 95% quantiles. Aspect ratio deviation increases with basal line tension. **e**, Distribution of cellular aspect ratio deviations for the experimental organoid in **b,c**. Dashed line is the median, dotted line perfect agreement (1.0) between experimental aspect ratio and theoretical expectation from the tensions. **f,g,h**, Median cell volumes, median cell heights and median aspect ratio deviations at different developmental time points with and without blebbistatin treatment. Conditions as in Fig. 3**i**. Cell volumes increase during budding, consistent with cell divisions. Aspect ratio deviations are lower than one for early stage organoids with a slight increase toward one at late times, consistent with the presence of small basal line tensions. As statistical test a Mann-Whitney-U-test was used. Black lines in violin plots indicate medians.

