## Supplementary Information for "Shape stabilization in budding intestinal organoids by geometrical ratcheting"

#### Contents

**Supplementary Figures** **3**

**Supplementary Tables** **19**

**Supplementary Video Captions** **22**

**Supplementary Note 1: Three-dimensional reconstruction method for organoids** **24**

**Supplementary Note 3: In-silico verifications of the force inference and pressure method in the two-cell and bubbly vertex model system, and for peaked distributions 39**

### Supplementary Figures

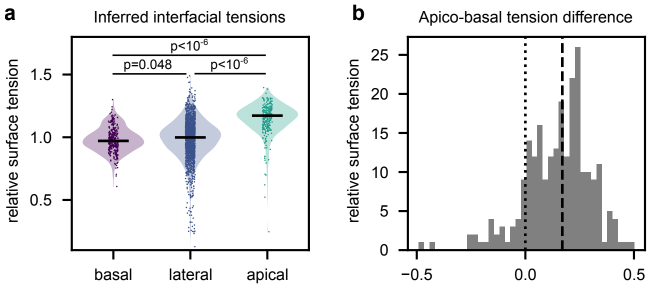

**Supplementary Figure S1 Surface tension distributions of budded organoid.** **a**, Distribution of inferred (relative) interfacial tensions by interface type for budded organoid 53 h post-splitting (cf. Fig. 3a). **b**, Distribution of single-cell apico-basal tension differences with zero marked as dotted line and median marked as dashed line.

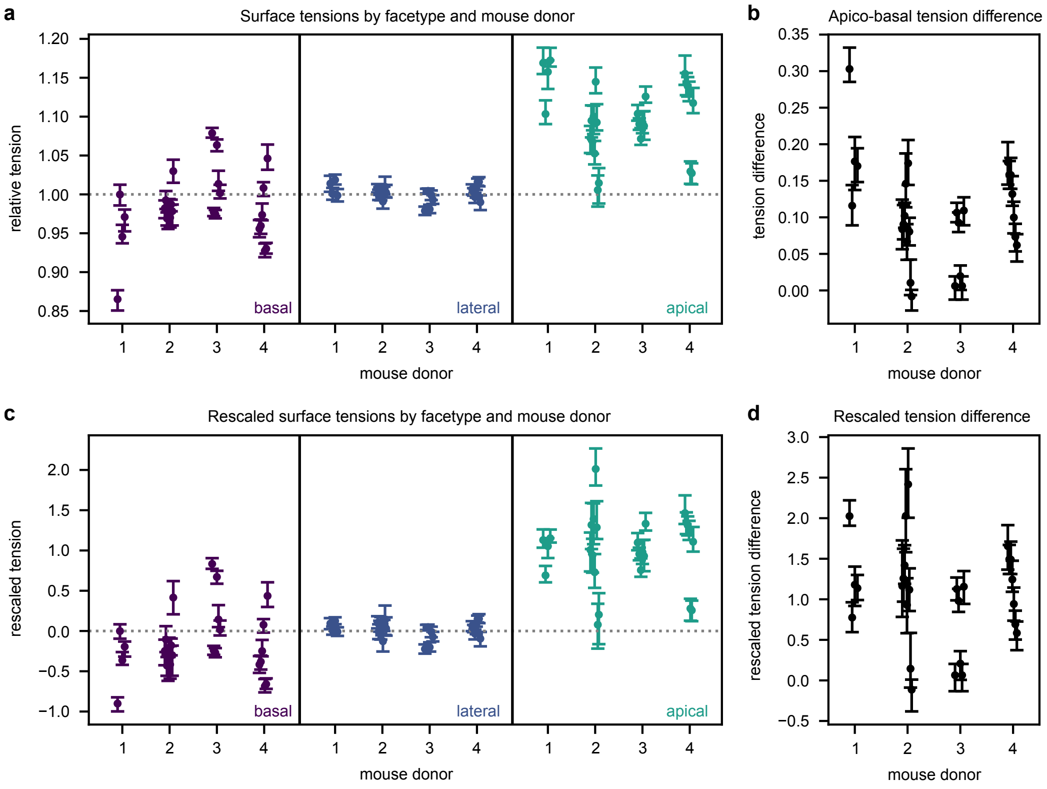

**Supplementary Figure S2 Interfacial tensions for different mouse donors. a**, Inferred relative interfacial tension medians by interface type with bootstrapped 95% confidence interval of the medians for individual 53 h control samples categorized by original mouse donor. **b**, Apico-basal tension difference medians with 95% confidence intervals for individual samples. **c,d** Interfacial tensions and apico-basal tension differences for individual samples after mouse-donor-specific normalization (see methods).

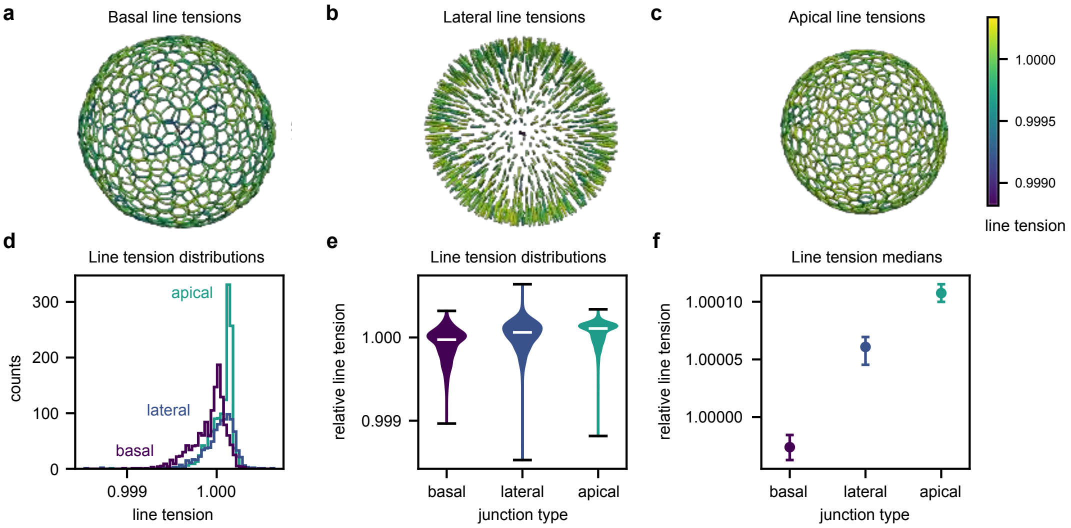

**Supplementary Figure S3 Junctional tangential force inference in unbudded in-silico organoid. a,b,c**, Reconstructed point cloud representation of tri-interfacial junctions, color-coded with the corresponding line tensions (tangential forces) for the unbudded organoid, cf. Fig. 1f. **d**, Histogram of inferred apical, basal and lateral tensions for the unbudded in-silico organoid subject to surface-tension based mechanics. **e**, Violin plots of the distributions for the different face types. Median is marked in white. **f**, Medians and bootstrapped 95% confidence intervals of the line tensions' medians.

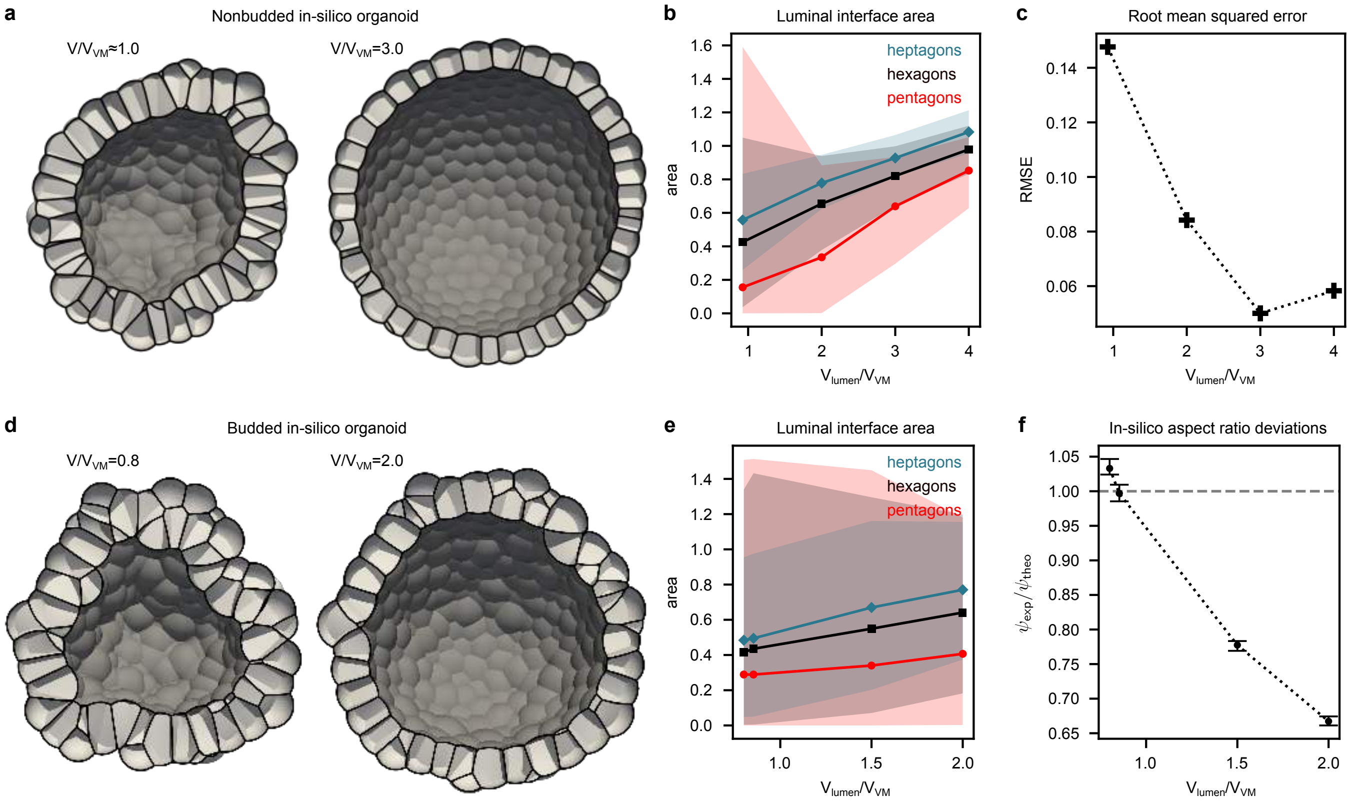

**Supplementary Figure S4 Effect of luminal volumes on simulated organoids.** **a**, Cross-section of unbudded in-silico organoid in Fig. 1f for different luminal volume ($V$) constraints, compared to the initial volume in the minimal energy configuration from the classical vertex model without interfacial curvature ($V_{\mathrm{VM}}$), where no volume constraint was assumed for $V/V_{\mathrm{VM}} \approx 1$. **b**, Luminal interfacial areas for different number of cell neighbors as function of the prescribed luminal volume. Interface sizes increase and collapsed interfaces are unfolded. Lines depict median with shaded area showing the range from 5% to 95% quantiles. **c**, Root mean squared error of the inference for different luminal volumes. The error decreases due to the growing luminal interface size. **d**, Cross-section of budded in-silico organoid in Fig. 5e for different luminal volume constraints. **e**, Luminal interface areas for budded organoid. **f**, Aspect ratio deviation for in-silico organoid for different luminal volume constraints. Increasing volume, i.e. increasing pressure, leads to a lowering of the observed aspect ratio due to mid-plane stretching. Errors bars indicate bootstrapped 95% confidence intervals around medians.

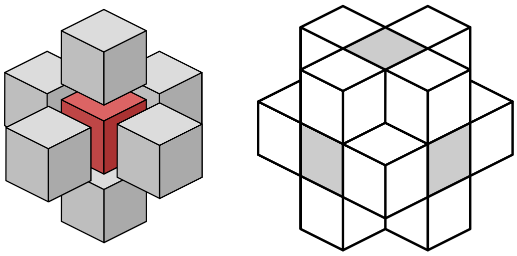

**Supplementary Figure S5 Schematic depiction of neighbor relations of voxels in 3D.** Around a voxel (red) the nearest neighbors form a cross (grey), corresponding to a 7-point 3D stencil (left). Going to the next-neighbor order yields additional voxels (white), in total forming three perpendicular neighbor planes, corresponding to the 19-point 3D stencil (right).

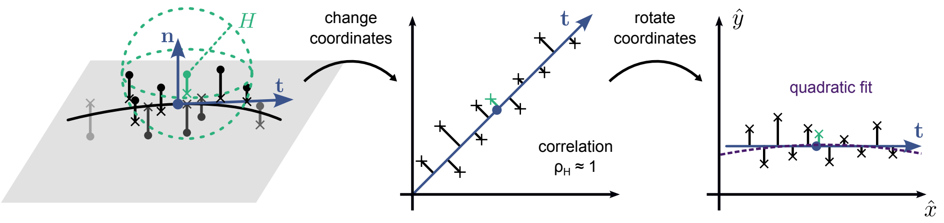

**Supplementary Figure S6 The curve thinning algorithm.** For a point cloud (left) the normal of the osculating plane $\mathbf{n}$ in a neighborhood with radius $H$ around point $\mathbf{x}_*$ (green) is determined via weighted principal component analysis (PCA) (blue). The points (circles) are projected onto the plane (crosses) and the tangent vector $\mathbf{t}$ is determined via weighted PCA. The tangent direction is then transformed onto the identity line (center) to assess the linearity within the neighborhood. For a linear curve the correlation $\rho_H$ is close to 1. The coordinates are then rotated (right) to perform a quadratic fit (dashed line). The centroid of the PCA is then considered as the projection of $\mathbf{x}_*$ onto the reconstruction. The centroid is projected onto the fit and is transformed back to the original coordinates.

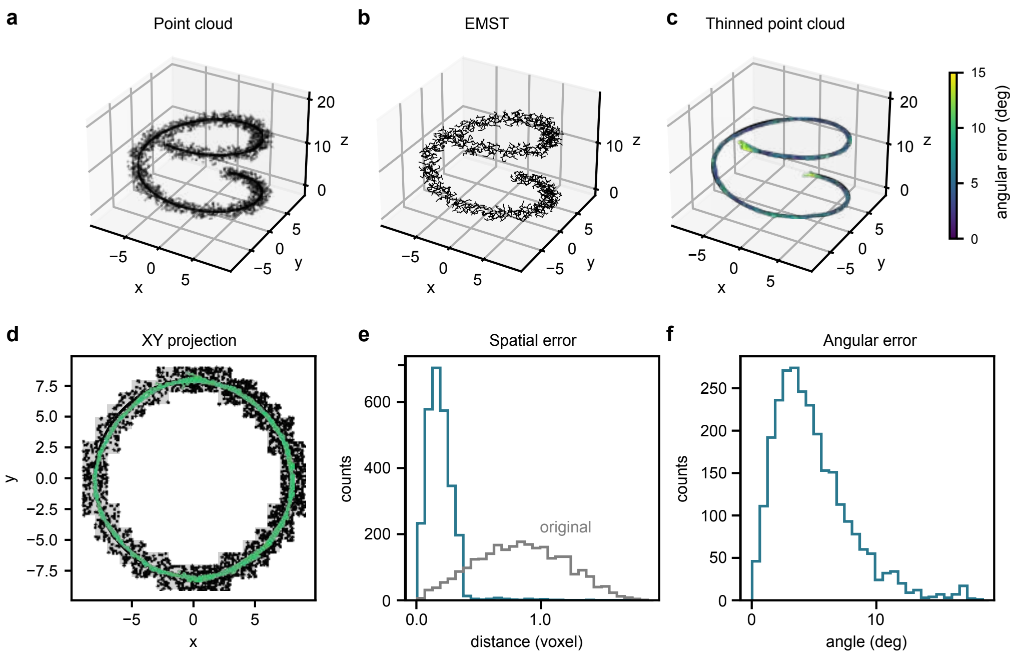

**Supplementary Figure S7 Curve thinning results for a known voxelated curve.** **a**, For a known curve (line) a voxelated volume and then a point cloud are created (circles). **b**, The euclidean minimum spanning tree (EMST) of the point cloud. **c**, The thinned point cloud, color coded with the angular error of the tangential vector. **d**, Projection of voxels (grey), original point cloud (black) and thinned point cloud (green) into the XY-plane. **e**, Spatial distance distribution of original and thinned points from curve. **f**, Angular error distribution for tangential vectors compared to theoretically expected tangents from curve.

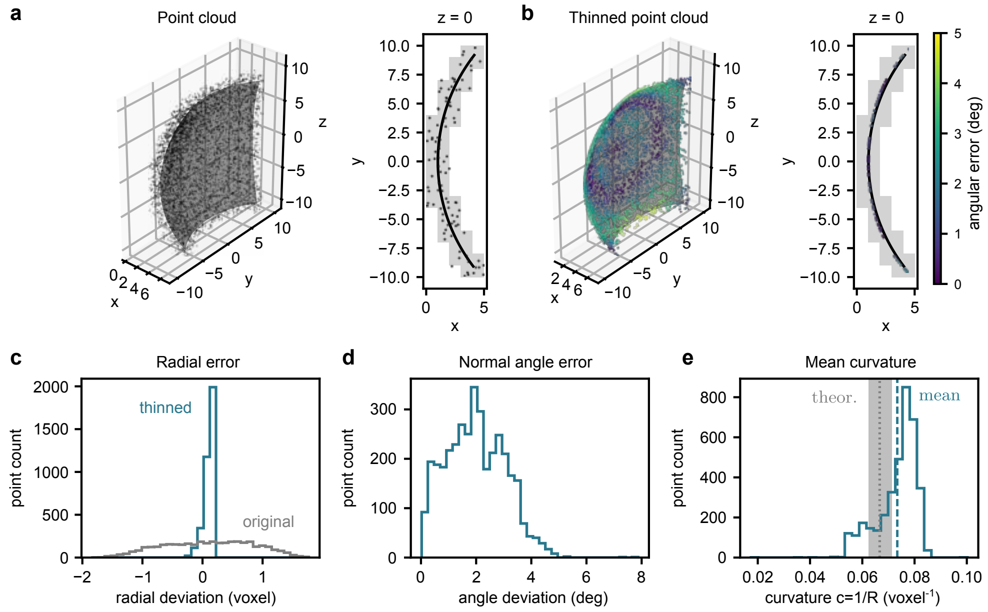

**Supplementary Figure S8 Interface thinning results for a known voxelated surface. a**, For a known spherical surface segment a voxelated volume and then a point cloud are created (circles). A cut through the $z = 0$ plane is shown by selecting the voxels from $z = 0$ to 1. **b**, The thinned point cloud, color coded with the angular error of the normal vector, again with the corresponding $z = 0$ cut. **c**, Histogram showing the distribution of the radial errors of the point clouds, i.e. the distance from known surface, for the original (grey) and thinned point clouds (blue). **d**, Distribution of the angular errors of the normal vectors after thinning and curvature reconstruction. Shown is the opening angle between the two normals. **e**, Mean curvature distribution from the reconstruction. For a sphere with radius $R = 15$ the mean curvature is $c = 1/R$ (dotted). The mean of the reconstruction (dashed) is compared to the interval corresponding to the interface thickness uncertainty $R \pm 1$ (grey).

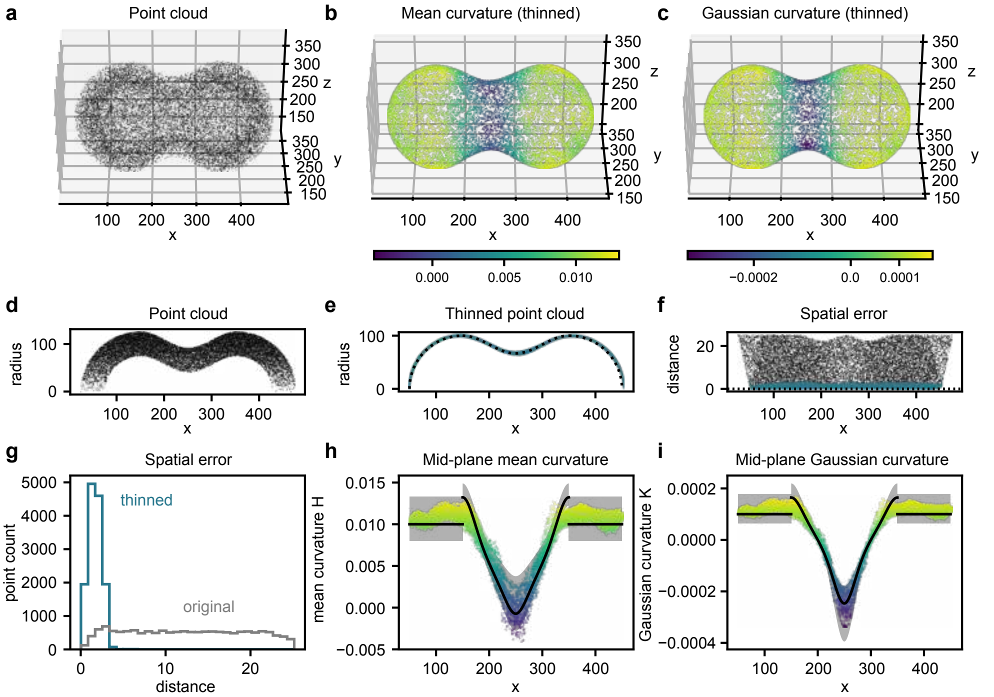

**Supplementary Figure S9 Organoid reconstruction results for a known voxelated dumbbell-shaped organoid. a**, For a known dumbbell-shaped organoid a voxelated volume and then a point cloud are created (circles). We consider a dumbbell surface with organoid thickness $\Delta r = 50$ voxels. **b,c**, The thinned point cloud, color coded with the mean curvature (**b**) and Gaussian curvature (**c**). **d,e**, The radial positions around the axis of symmetry of the points representing the organoid as a function of the position along the axis for both the original point cloud (**d**) and the thinned one (**e**). The dotted line is the known mid-plane radius. **f**, The spatial error, i.e. distance from the surface, of both the original (black) and the thinned (blue) point cloud along the axis of symmetry. **g**, The distribution of the spatial errors for both the original (grey) and thinned (blue) point clouds. **h,i**, The reconstructed mean (h) and Gaussian curvatures (i) of the points along the axis of symmetry, color-coded with the corresponding values as in **b**,**c**. The theoretical curvatures are shown as a line with the uncertainty from the organoid thickness in grey, given by the curvatures of the inner surface at radius $r - \Delta r/2$ and the outer surface at $r + \Delta r/2$.

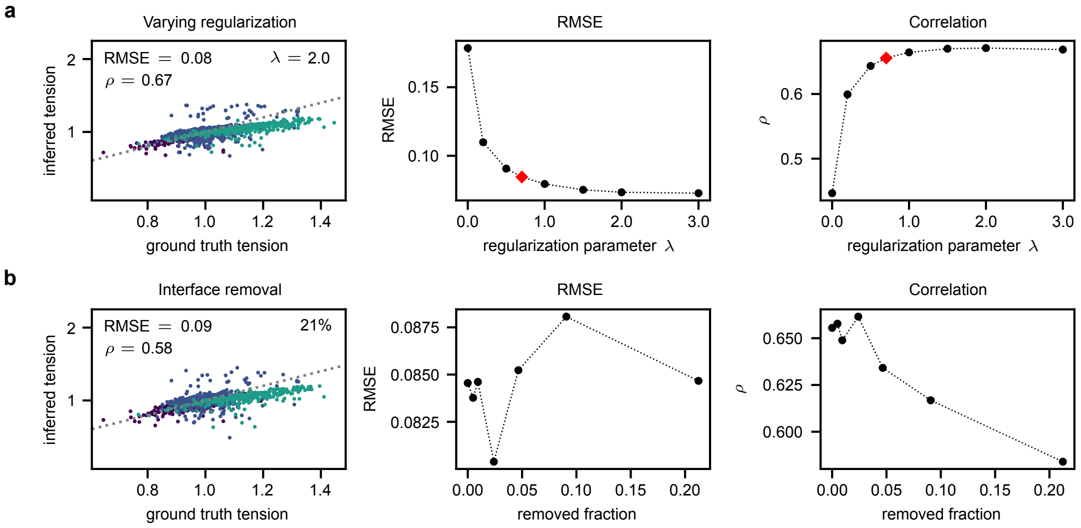

**Supplementary Figure S10 Robustness of force inference.** **a**, To test robustness against the choice of the regularization parameter $\lambda$, different results for the in-silico organoid from Fig. E2 with different levels of regularization are shown. Root mean squared error (RMSE) decreases and correlation $\rho$ increases with larger $\lambda$, but for large $\lambda$ the slope of inferred tension does not match the ground truth data. The red marker is the optimal result with respect to the PRESS statistics. **b**, The robustness with respect to random interface removal is shown. The RMSE and correlation are robust even for a removal of approximately 10% of all interfaces.

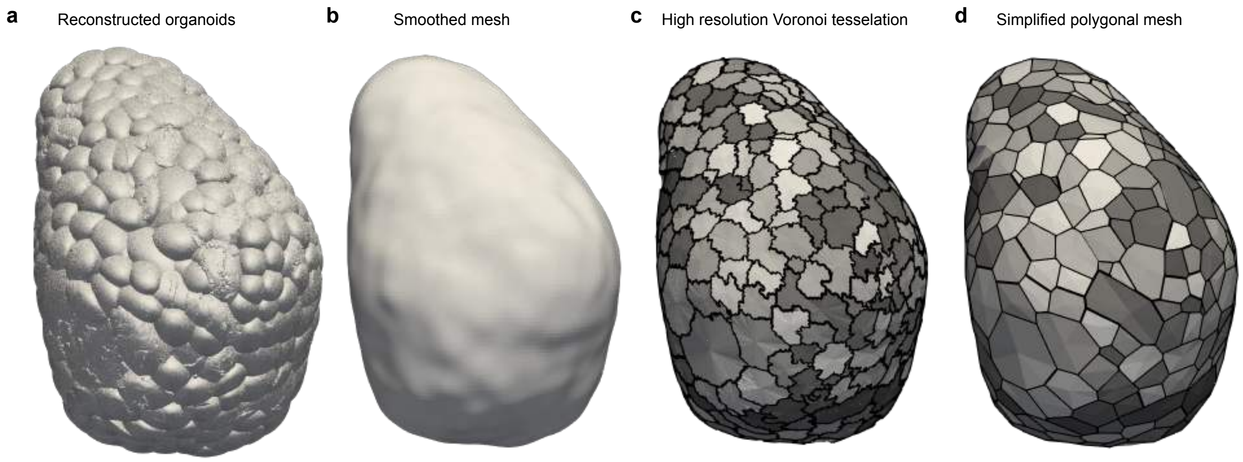

**Supplementary Figure S11 Vertex model mesh extraction from point-cloud based reconstruction.** **a**, The three-dimensional reconstruction is used as a starting point for vertex model (VM) mesh extraction. **b**, From a binary mask of the organoid volume a smoothed mesh is created to whose surface the points from the point cloud are projected. **c**, For these projected points a Voronoi-like tesselation on a dual mesh is created and assigned to the individual interfaces of the organoid, using the assignment from the point cloud (color-coded). This yields a high-resolution Voronoi-like mesh. **c**, The high-resolution mesh is simplified by only keeping interfacial boundaries as straight lines and points where three separate interfaces meet.

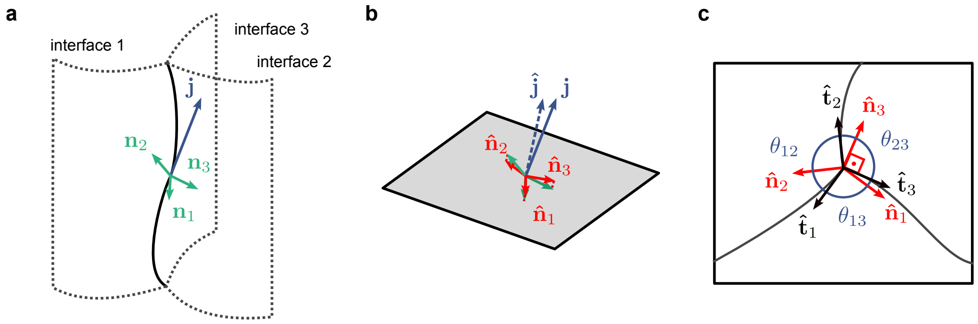

**Supplementary Figure S12 Schematic depiction of dihedral angle reconstruction along juction. a**, Along a junction with tangential vector $\mathbf{j}$ the corresponding interfaces can be described by normal vectors $\mathbf{n}_i$ for $i = 1, 2, 3$. **b**, As the interface normals should lie in the orthogonal space of the tangent, we may estimate the junctional tangental vector $\hat{\mathbf{j}}$ by considering the averaged cross-products of the normal vectors. The normal vectors are then projected onto the space perpendicular to the averaged tangential $\hat{\mathbf{j}}$, reading $\hat{\mathbf{n}}_i$. **c**, In this perpendicular space the projected normals $\hat{\mathbf{n}}_i$ are orthogonal to the (projected) interface tangents $\hat{\mathbf{t}}_i$, allowing the calculation of the dihedral angles $\theta_{ij}$ at this junctional point via the normals.

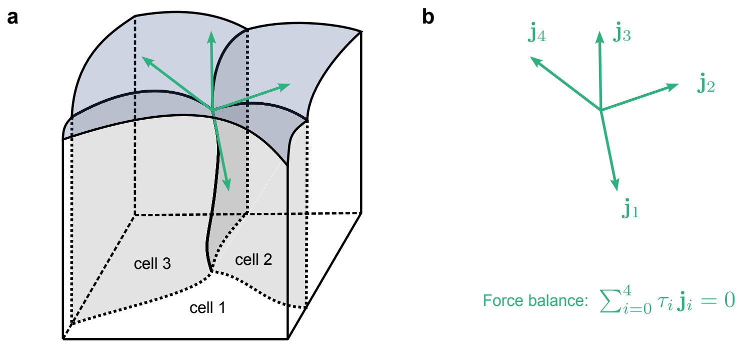

**Supplementary Figure S13 Schematic depiction of tangential force inference. a**, For multiple junctions between interfaces meeting at a point (junctional vertex) the tangents point in different directions (green). **b**, Assuming mechanical equilibrium the tangential forces with magnitude $\tau_i$ along the junctions must cancel out at junctional vertices due to force balance.

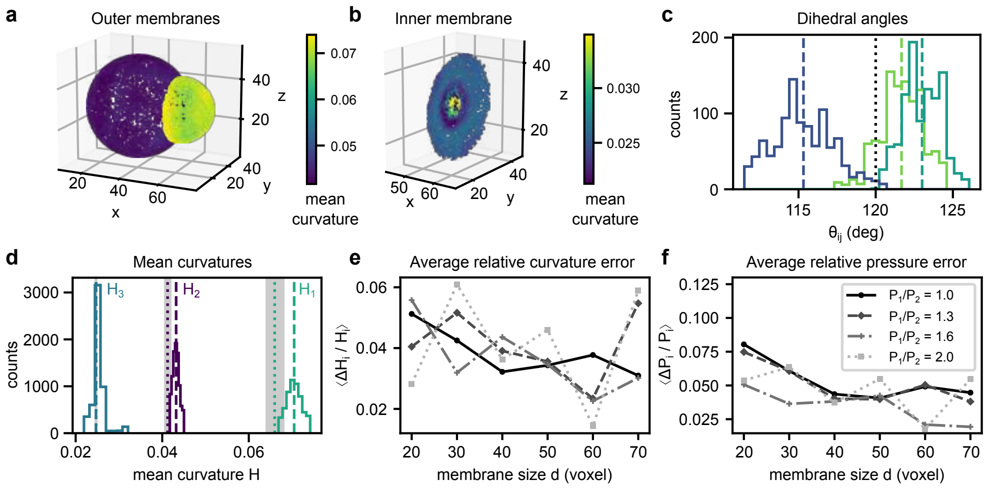

**Supplementary Figure S14 Pressure inference results for a prescribed in-silico two-cell system.** For a two cell system (cf. Extended Data Fig. 1a) with prescribed pressures and identical interface tensions $\gamma_i = 1$, the interfaces are reconstructed and the pressures inferred. **a,b**, Reconstructed interfaces color-coded with the mean curvature calculated at the reconstructed points. **c**, Distribution of the reconstructed dihedral angles $\theta_{ij}$. Due to the identical tensions the theoretical angle is 120°, depicted as black dotted line. **d**, Distributions of the mean curvatures for the different interfaces. Outliers outside the region from the 0.01 to the 0.99 quantiles are not shown. The reconstructed average mean curvatures (including outliers, dashed) are close to the theoretical curvatures (dotted) but outside the radial error margin from voxelation $R \pm 0.5$. **e,f**, For different resolutions, corresponding to interface sizes $d$, the errors in the pressure inference determined. Different pressure ratios $P_1/P_2$ in color. The average relative curvature error (with respect to the largest curvature $H_1$) (**e**) and the average relative pressure errors (**f**) are shown.

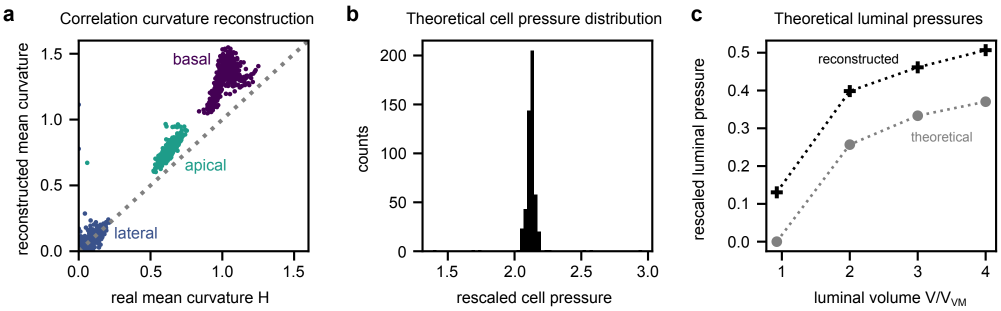

**Supplementary Figure S15 Verification of cellular and luminal pressure inference in the bubbly vertex model.** **a**, For an organoid simulated with the bubbly vertex model real mean curvatures, computed via Surface Evolver [1], and the reconstructed mean curvatures correlate. Outliers in real and reconstructed curvatures from collapsed interfaces are not shown (4 interfaces). Dashed line is the identity. **b**, Cellular pressures, indicated as pressure difference to the background pressure, show a sharp distribution at $P \approx 2$ in dimensionless units, where some single cells deviate (expanding the $x$-axis). **c**. Theoretical (grey) and inferred (black) luminal pressures as a function of (prescribed) lumen volume $V/V_{\mathrm{VM}}$ with initial vertex model volumes $V_{\mathrm{VM}}$. The theoretical pressure is obtained from the Lagrange multiplier in the simulation. The first point at $V/V_{\mathrm{VM}} \approx 1$ corresponds to the theoretically unpressurized case, $P_{\mathrm{lumen}} = 0$. The volume $V/V_{\mathrm{VM}} = 3$ corresponds to **b,c**.

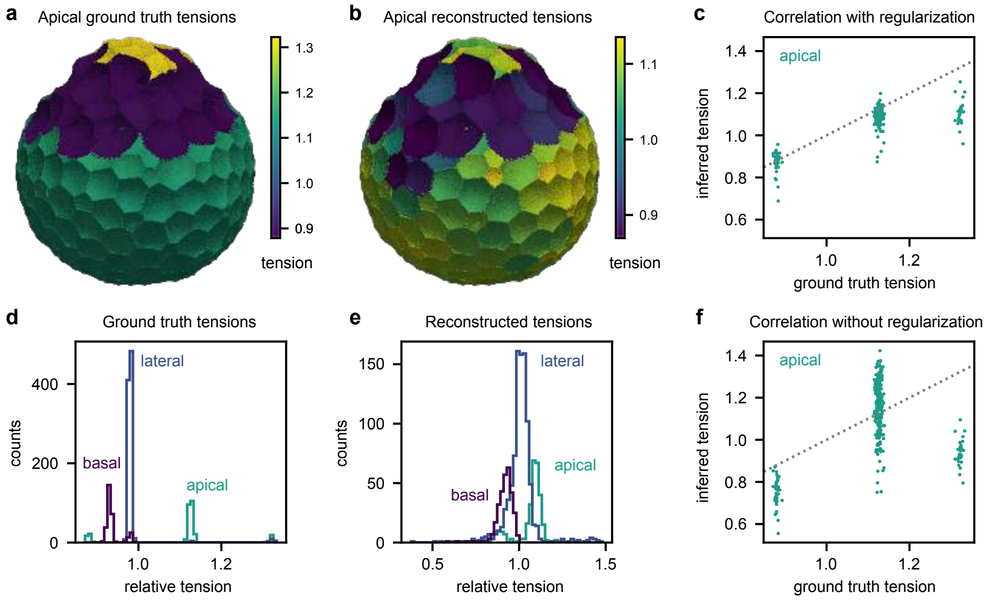

**Supplementary Figure S16 Force inference of a budded pressurized in-silico organoid with strongly peaked tension distributions.** **a**, Reconstructed organoid with apical ground truth tensions (color-coded with range between 5% and 95% quantiles, top) and distribution of tensions (bottom). The organoid contains 300 cells, has a luminal volume of $V/V_{\mathrm{VM}} = 2$, and has strongly peaked distributions of tensions. **b**, Inferred (apical) tensions (top) with distribution of tensions (bottom) with Thikonov regularization. **c,d**, Correlation of inferred apical tensions and real ground truth tensions with (**c**) and without Thikonov regularization (**d**). The distributions from which the tensions are drawn are Gaussian with a standard deviation of 0.005 and means of 1 for lateral, 1.15 and 1.35 for apical, and 0.95 (buds) and 1.0 for basal faces.

### Supplementary Tables

| Donor | Replicate | 5h ctrl | 48h ctrl | 53h ctrl | 5h bleb | 48h bleb |
|---|---|---|---|---|---|---|
| 1 | – | – | – | 4 | 5 | – |
| 2 | R1 | 2 | 4 | 2 | 7 | 4 |
| 2 | R2 | 4 | 2 | 4 | 6 | 1 |
| 2 | R3 | – | 6 | 3 | 1 | 2 |
| **2** | **sum** | **6** | **12** | **9** | **14** | **7** |
| 3 | R1 | – | – | 1 | 1 | 1 |
| 3 | R2 | 2 | – | 1 | – | 4 |
| 3 | R3 | 2 | 2 | 4 | 2 | 3 |
| **3** | **sum** | **4** | **2** | **6** | **3** | **8** |
| 4 | R1 | 3 | 3 | 4 | 6 | 3 |
| 4 | R2 | 2 | 1 | 1 | 2 | 4 |
| 4 | R3 | 5 | 2 | 2 | 3 | 3 |
| **4** | **sum** | **10** | **6** | **7** | **11** | **10** |
| Total (all donors) | | 20 | 20 | 26 | 33 | 25 |
| Total (excl. donor 1) | | 20 | 20 | 22 | 28 | 25 |

**Supplementary Table S1** Details on donor and replicate sample sizes. Donor 1 is only used for the comparison of control samples at 53 h with the blebbistatin-treated samples (B5 h).

| Name | Company | Catalog number | Dilution | Final concentration |
|---|---|---|---|---|
| M-anti-$\beta$-Catenin | BD Transduction Laboratories | 610154 | 1:500 | |
| Rb-anti-Lysozyme | DAKO | A0099 | 1:300 | |
| Goat-anti-Mouse-AF488 | | A11011 | 1:500 | |
| Donkey-anti-Rabbit-647 | | A32795 | 1:500 | |
| Hoechst | Thermo Scientific | 62249 | | 1 $\mu$g/ml |
| Phalloidin-TRITC | Sigma | P1951 | | 50 ng/$\mu$l |

**Supplementary Table S2** Antibodies and dyes used in this study.

| | First surface thinning | Second surface thinning | Curvature calculation |
|---|---|---|---|
| $H_0$ | $1.5\,t$ | $0.75\,t$ | $0.6\,t$ |
| $\Delta H$ | $2\ \max\{\Delta x, \Delta y, \Delta z\}$ | $2\ \max\{\Delta x, \Delta y, \Delta z\}$ | $2\ \max\{\Delta x, \Delta y, \Delta z\}$ |
| $\hat{H}$ | $5\,H_0$ | $5\,H_0$ | – |
| $d_{\mathrm{cutoff}}$ | $5\ \max\{\Delta x, \Delta y, \Delta z\}$ | $5\ \max\{\Delta x, \Delta y, \Delta z\}$ | $5\ \max\{\Delta x, \Delta y, \Delta z\}$ |

**Supplementary Table S3** Reconstruction hyperparameters for organoid geometry reconstruction.

### Supplementary Video Captions

***Supplementary Video 1: Organoid microscopy data and segmentation.***

Three-dimensional volume renderings of the microscopy data of a sample budded intestinal organoid (left) and the corresponding voxel-wise segmentation (right). In microscopy data green is membrane (beta-catenin), blue is nucleus and red is F-actin, cf. Fig. 1c.

***Supplementary Video 2: Three-dimensional surface tensions for budded intestinal organoid.***

A three-dimensional rendering of the reconstructed interfaces of a sample budded intestinal organoid for basal (left), lateral (center) and apical (right) interfaces. Interfaces are reconstructed as point clouds, where individual points are depicted as small spheres. The data is color-coded by the relative interfacial tension with the color map depicting the tensions from the 5% to the 95% quantiles, cf. Fig. 3a.

***Supplementary Video 3: Splitting of interfacial tensions into budded regions and non-budded regions.***

A three-dimensional rendering of the reconstructed interfaces of a sample budded intestinal organoid for basal (left), lateral (center) and apical (right) interfaces. Interfaces are reconstructed as point clouds, where individual points are depicted as small spheres. Buds in the organoid are selected manually in microscopy data as spherical regions (black) to split interfaces into subpopulations inside and outside the buds. The data is color-coded by the relative interfacial tension with the color map depicting the tensions from the 5% to the 95% quantiles.

***Supplementary Video 4: Inferred mid-plane curvatures for budded intestinal organoid.***

A three-dimensional rendering of the reconstructed tissue mid-plane of a sample budded intestinal organoid. The surface is reconstructed as a point cloud, where individual points are depicted as small spheres. The data is color-coded by the mean curvature (left) depicting the values from the 5% to the 95% quantiles and by the Gaussian curvature (right), with color boundaries set as such that the sign can be seen, cf. Fig. 3f.

***Supplementary Video 5: Ground truth interfacial tensions and reconstructed mid-plane for in-silico organoid.***

A three-dimensional rendering of a reconstructed budded in-silico organoid (top), color coded by the relative ground truth surface tensions (from 5% to 95% quantiles) for basal (left), lateral (center) and apical (right) interfaces. The mid-plane curvatures are inferred (bottom) and shown with color-coding from 5% to 95% quantile in mean curvature (bottom left) and with color-coding showing the sign in Gaussian curvature (bottom right).

***Supplementary Video 6: Vertex model simulations of growing fluidized organoids.***

A three-dimensional (flat interface) vertex model simulation of a growing fluidized organoid with three different fluidization levels, expressed as the rate of topological T1 transitions per cell $\nu$, cf. Fig. 4a. Shown is the shape in simulation time. Color-coded are the different cell types, i.e. stem cells, transit amplifying (TA) cells, and differentiated cells.

***Supplementary Video 7: Simulated organoid relaxation upon fluidization for reconstructed mouse small intestine organoids.***

In-silico relaxation of experimentally observed and computationally reconstructed organoids with the (flat interface) vertex model, cf. Fig. 4j. Shown are three different organoids with different complexities relaxed over dimensionless simulation time. The rate of topological T1 transitions per cell was chosen as $\nu = 2$. Only inferred topologies are used for relaxation, i.e. cell volumes are assumed to be $V = 1.0$, and no apico-basal asymmetry is assumed ($\Gamma_\mathrm{a} = \Gamma_\mathrm{b} = \Gamma_\mathrm{l} = 1.0$) for the relaxation.

***Supplementary Video 8: Three-dimensional line tensions for budded intestinal organoid.***

A three-dimensional rendering of the reconstructed tri-interfacial junctions of a sample budded intestinal organoid for basal (left), lateral (center) and apical (right) junctions. Junctions are reconstructed as point clouds, where individual points are depicted as small spheres. The data is color-coded by the relative line tension with the color map depicting the tensions from the 5% to the 95% quantiles, cf. Fig. 5a.

### Supplementary Note 1: Three-dimensional reconstruction method for organoids

#### Three-dimensional segmentation of organoid microscopy data

In order to estimate dihedral angles between the interfaces between cells or between a cell and their surroundings from microscopy data, we first need to obtain a three-dimensional representation of the cells. From the fluorescence microscopy data we obtain a three-dimensional grid of fluorescence intensities, which yields three-dimensional voxel-wise information, with voxels being volume pixels. As a first step of the analysis we generate a segmentation, where cell identities are assigned to each voxel to obtain a representation of the cells on the imaging grid. Image segmentation is a classical problem in computer vision, and has been investigated thoroughly in the context of biological cells.

The microscopy data, which is recorded with a 25x objective and a pixel size of $250\,\mathrm{nm} \times 250\,\mathrm{nm}$ at a slice-to-slice distance of 500 nm, is cropped and interleaved by introducing an averaged image slice between two recorded slices. This yields cubic voxels to be analyzed. We observed an intensity loss as we imaged slices further away from the glass slide, due to light scattering and absorption in tissue through which we have to image. To improve the results a tissue clearing technique was used, reducing scattering effects in the imaging of large samples [2], and laser intensity increased throughout the organoid. However, we still see an intensity loss. To adjust for this, we perform a linear intensity correction, where the pixel values in individual slices are scaled linearly to yield a visually uniform imaging intensity. Further preprocessing includes image registration [3], i.e. the images are rotated and translated to correct for sample movement during image acquisition, and gamma correction, i.e., pixel values are processed with a power law function to increase intensity in dark pixels, if necessary for the individual organoid data. The organoids were chemically fixed, i.e. their shape was stationary. Then antibody staining was performed, where $\beta$-catenin (green), primarily situated in the plasma membrane, filamentous actin (magenta), enriched on the apical cell side, and DNA (blue) in the nucleus were fluorescently labeled. The interfaces are thus primarily determined through the green and red channels, which were selected as a boundary image.

This boundary image now serves as the starting point for voxel segmentation. We use Cellpose 2.0 [4, 5], which is a tool leveraging artificial intelligence via a convolutional U-net for microscopy image segmentation. Through a man-in-the-loop training procedure we trained a specialized network for three-dimensional organoid image stacks by adding manually corrected segmentations from Cellpose to the training data. The "cyto" Cellpose model was used as a starting model, which was fine-tuned using our annotated data. The training samples included perpendicular sections through preprocessed organoid image stacks from three sample organoids, which were not used further within the study. Our trained Cellpose model now yields a voxel-wise segmentation of the microscopy data, cf. Cellpose mask in Fig. 1d. However, we found that Cellpose tends to oversegment the volumes, as it performs two-dimensional segmentation in the three perpendicular directions and combines these in a last step to obtain three-dimensional segmentations. The algorithm yields reasonable cell boundaries, introduces, however, too many such boundaries.

To improve upon this result, we then use ilastik [6], which uses classical machine learning approaches for segmentation, but allows for the native use of three-dimensional image processing filters. As input for ilastik we use the data including the nucleus channel and an edge probability map. To create the latter we take the Cellpose segmentation, use an edge filter on the data [7], which assigns a probability of 1 to boundary voxels of cells, and then perform Gaussian blurring. This yields a smoothed probability map for edges, cf. Fig. 1d. Based on this input data ilastik creates an oversegmented segmentation using a watershed algorithm, and trains a classifier on resulting edges, to merge areas into the final segmentation. Due to the three-dimensional nature of the used filters this method takes advantage of the three-dimensional structure of the data and we found it to perform better at obtaining a good three-dimensional segmentation. However, three-dimensional segmentation with this workflow is still not perfect and yields some cells containing only few voxels. In order to make reliable deductions from experiments, we therefore perform post-processing in which we leave out cells suffering from wrongly segmented interfaces. In addition, wrongly segmented cells with too few voxels (here 1000 voxels) are ignored in the analysis. This is justified due to the overdetermined nature of the inference problem.

#### Reconstruction of tri-interfacial junction curves

As starting point we consider the cellular segmentation. Each voxel which has three different cells as next-nearest-neighbors (corresponding to a 19-point stencil in the 3D lattice, cf. Supplementary Fig. S5) is considered a junctional voxel.

For each junction, representing the curve of three touching cells and interfaces, all voxels are considered and transformed into a point cloud representation. For this we consider a 3D rectangular voxel $V_{(x,y,z)} = [x, x + \Delta x] \times [y, y + \Delta y] \times [z, z + \Delta x]$ with lattice spacings $\Delta x, \Delta y, \Delta z$ in $x$, $y$ and $z$-directions, respectively. Each of these voxels is subsampled by drawing $N_{\text{sub}}$ points $\mathbf{x}_1, \dots, \mathbf{x}_{N_{\text{sub}}} \in V_{(x,y,z)}$ from the uniform distribution $U\left(V_{(x,y,z)}\right)$.

##### Curve thinning and tangent estimation

Taking all subsampled points from all junctional voxels yields a point cloud representation of the junction. The point cloud now has a finite thickness due to the choice of the stencil and the image resolution. The real junction is ideally a one-dimensional curve in space, where the point cloud can be interpreted as a sample drawn from a distribution centered around the true curve. As such we adapt a curve thinning approach to infer the true curve non-parametrically introduced by Lee [8], which we explain in the following.

First, the euclidean minimum spanning tree (EMST) is constructed. The EMST is a tree, i.e. a graph in which there is a unique path from every point to another point, which minimizes the total edge length of the edges connecting the points. We consider a special point $\mathbf{x}_* \in \{x_i\}$, i.e. a specific point from the point cloud that we want to project onto an optimal local curve which we infer from the point cloud in a neighborhood with radius $H$. The algorithm that we now introduce is depicted schematically in Supplementary Fig. S6. As we do not want mixing with points far away along the curve, even if they are within radius $H$, we construct the neighborhood by propagating along the EMST, starting from $\mathbf{x}_*$, adding points to the neighborhood until the points' distances from $\mathbf{x}_*$ exceed $H$. On this neighborhood an osculating plane is calculated via weighted principal component analysis (PCA) [9]. We consider the weight for point $\mathbf{x}_i$ [8]

$$w_i = \begin{cases} 2\frac{\|\mathbf{x}_* - \mathbf{x}_i\|^3}{H^3} - 3\frac{\|\mathbf{x}_* - \mathbf{x}_i\|^2}{H^2} + 1, & \text{if } \|\mathbf{x}_* - \mathbf{x}_i\| < H \\ 0, & \text{else} \end{cases}. \tag{S1}$$

With the weight function we account for the fact that points close to $\mathbf{x}_*$ are more relevant when determining the (local) linear regime and should thus be considered more strongly.

The weighted centroid is now computed via

$$\mathbf{c} = \sum_{i=1}^{N_{\text{sub}}} \frac{w_i \mathbf{x_i}}{\sum_{j=1}^{N_{\text{sub}}} w_j} \tag{S2}$$

and with weight matrix $\mathbf{W} = \text{diag}(w_1, \ldots, w_{N_{\text{sub}}})$ and distance matrix $\mathbf{Y} = (\mathbf{x}_1 - \mathbf{c}, \ldots, \mathbf{x}_{N_{\text{sub}}} - \mathbf{c})^T$ we determine the principal directions as the eigenvectors of the weighted scatter matrix $\mathbf{Y}^T\mathbf{W}\mathbf{Y}$. In the direction of the eigenvector corresponding to the smallest eigenvalue the point cloud shows the least variance, meaning this (normalized) eigenvector represents the normal $\mathbf{n}$ of the osculating plane.

This can be seen by considering the analogous observation that the normal $\mathbf{n}$ and the (unweighted) centroid $\mathbf{c}$ minimize the squared distance from a (2D) hyperplane, i.e. they solve

$$\min_{\mathbf{c},\mathbf{n},\|\mathbf{n}\|=1} \sum_{i=1}^{N_{\text{sub}}} \left((\mathbf{x}_i - \mathbf{c}) \cdot \mathbf{n}\right)^2 . \tag{S3}$$

Next, the points in the neighborhood are projected onto the osculating plane by reexpressing the distances $(\mathbf{x}_i - \mathbf{c})$ in the normalized principal components and neglecting the component normal to the plane. For the projected points again a weighted PCA is performed to find the normal (small eigenvalue) and the tangent (large eigenvalue) of the curve in 2D. To quantify how linear the curve is in the chosen neighborhood, the coordinate system can be rotated as such that the tangent now points in the direction of the identity function, i.e. in $(1, 1)$-direction. For these transformed points we can determine the quality of a linear fit by calculating the Pearson coefficient of correlation (for given radius $H$) $\rho_H$ [8]. For a perfectly linear curve, the points now fall onto the identity function and the correlation is maximal $\rho_H = 1$.

Assuming $H$ is chosen such that the curve is in a linear regime, we perform a quadratic regression. For this the points are rotated in the osculating plane such that the normal of the curve (perpendicular to the tangent) points into the $\hat{y}$-direction. The points $(\hat{x}_i, \hat{y}_i)$ in this coordinate system are then fitted with a quadratic function.

We consider the vector $\mathbf{y} = (y_1, \ldots, y_{N_{\rm sub}})^T$, design matrix

$$\mathbf{X} = \begin{pmatrix} \hat{x}_1^2 & \hat{x}_1 & 1 \\ \vdots & \vdots & \vdots \\ \hat{x}_{N_{\rm sub}}^2 & \hat{x}_{N_{\rm sub}} & 1 \end{pmatrix}, \tag{S4}$$

and the weight matrix (recomputed in the new coordinate system) $\mathbf{W}$ to determine the coefficients of the quadratic fit via weighted least squared error (WLSE) estimate to be [8, 10]

$$\hat{\beta} = \left(\mathbf{X}^T \mathbf{W} \mathbf{X}\right)^{-1} \mathbf{X}^T \mathbf{W} \mathbf{y}. \tag{S5}$$

The coefficient vector $\hat{\beta} = (\beta_2, \beta_1, \beta_0)$ contains the quadratic, linear and constant terms in the regression $\beta_2$, $\beta_1$ and $\beta_0$, respectively, i.e. $\hat{y} = \beta_2 \hat{x}^2 + \beta_1 \hat{x} + \beta_0$. In this coordinate system the original point $\mathbf{x}_*$, around which the neighborhood is centered, is projected onto the centroid, corresponding to the point $(0, \beta_0)$ on the regression curve. By transforming back into the original coordinates we project the point onto the underlying curve, leading to a thinned curve. The tangent of the curve at the particular point was estimated in the second PCA step. Note, that it is also possible to estimate the tangent of the intersection of three interfaces from interface normals alone [11], which we use in our force inference method, see Supplementary Note 2.

Note that in the original algorithm by Lee [8] a WLSE method with a predefined coordinate system is chosen instead of the described weighted PCA steps. The PCA error on the normals and tangents and the PCA method itself are, however, independent of the choice of the coordinate system. This is due to the minimization of the variance orhthogonal to the plane/line contrary to WLSE, where the variance is with respect to the (arbitrarily chosen) $z$-direction, cf. Supplementary Fig. S6. As such it is not biased from the choice of a coordinate system.

This curve thinning algorithm has been found to be more robust than a moving average approach [8], as considered by Xu et al. [11]. Indeed, we also find a better junctional reconstruction because moving averages will be biased toward the side to which the curve bends, as the density of points is larger there.

#### Neighborhood size optimization

The curve thinning algorithm assumes a neighborhood radius $H$ which captures the linear regime around the special point $\mathbf{x}_*$ that is being projected onto the curve and then considers quadratic contributions in the fitting step. The linearity within the chosen neighborhood can be quantified considering the correlation $\rho_H$ after coordinate transformation. In the original approach $H$ is initialized as slightly larger than the estimated thickness of the point cloud and then successively increased until the correlation exceeds a cutoff criterion [8].

The optimal $H$ will depend on both point cloud thickness and curvature. For our purposes the point cloud thickness is determined by the choice of the 19-point stencil and the curvature along the junctions does not vary profoundly. As such we choose a representative point (with an index in the middle of the point cloud) and increase $H$ staring from 4 voxels in steps of 0.1 voxels until $\rho_H > 0.9$ or until $H > 20$, taking the $H$ with maximum correlation in the latter case. Note that we perform the curve reconstruction in real space and not in voxel space, i.e. we multiply the voxel positions by $\Delta x$, $\Delta y$, and $\Delta z$ for the respective coordinates. The neighborhood radius $H$ is determined independently for all junctions but within the junction the found $H$ will be used for all points in the thinning algorithm.

#### Comparing reconstruction results to ground truth data

To verify our curve thinning algorithm we generate data which is of similar nature as the junctional data obtained from organoid segmentations and test our approach on this data. For this we consider the curve

$$\mathbf{r}(u) = \begin{pmatrix} r\,\sin(u) \\ r\,\cos(u) \\ 2\,u \end{pmatrix}, \quad \text{for } u \in [0, 3\pi], \tag{S6}$$

with radius $r = 8$ (in voxel space). As the determination of junctional voxels is done via the 19-point stencil in 3D (cf. Supplementary Fig. S5) a segmented curve will consist of multiple voxels in the tangential direction, i.e. have a finite thickness. To recreate this we construct a curve with a finite radial thickness of $\delta r = 0.5$. By considering points along $r(u)$ and drawing additional points from a uniform distribution inside a sphere around the points on the curve we sample the volume of the curve. We consider the curve in voxel space (with lattice constant 1) and determine the voxels through which the curve runs by considering the sampled points. These voxels are then treated like the voxels from the segmentation. They are subsampled with 8 points. In organoids we subsample with 6 points, which yields similar results as we have more voxels from the consideration of the 19-point stencil. Supplementary Fig. S7 shows the data and the thinning results. The point cloud around the true curve is used for the thinning algorithm, cf. Supplementary Fig. S7a. The EMST corresponding to the point cloud, used for neighbor determination, is shown in Supplementary Fig. S7b. The thinned point cloud indeed lies on the curve, Eq. (S6), with small deviations at the two ends of the curve, cf. Supplementary Fig. S7c. In the projection of the data onto the $xy$-plane (Supplementary Fig. S7d), this can be seen more clearly with the thinned point cloud lying within the voxels and very close to the original curve except at the ends, situated at the outer most points along the $y$-axis. To quantify this, the histograms of the spatial error, i.e. the distance of the points from the original curve, of both the original and the thinned curve are shown in Supplementary Fig. S7e with a clear decrease of the error into the regime of below half a voxel. The angular error of the tangential vectors was determined for the tangents at the thinned points and we find an angular error of the order of $0°$ to $10°$, see the histogram Supplementary Fig. S7f, with larger errors at the end of the curve (Supplementary Fig. S7c).

While the thinning works as expected and leads to good reconstruction results without any apparent spatial bias (opposed to a moving averages approach), we find non-negligible errors for the tangential vectors. For this reason we adopt a different tangential vector estimating technique, solely using the interfaces' normal vectors in the force inference algorithm.

#### Surface reconstruction

Similarly to the reconstruction of the tri-interfacial junctions, we start with the cell segmentation. Now, each voxel which has two different cells as next-nearest-neighbors (corresponding again to a 19-point stencil, cf. Supplementary Fig. S5) is considered an interface voxel. Additionally, we consider the junctional voxels which contain the same two cells and add them as additional voxels to the interface, meaning these junctional voxels are used for three interface reconstructions. In the same manner as for the junctions, point clouds are created via subsampling. For our purposes the reconstruction must yield surface normals for angle calculations at junctions and surface curvatures. To do this, we again follow the point cloud based approach suggested in Ref. [11] and generalize the curve thinning algorithm by Lee [8] to a surface thinning algorithm.

##### Initial normal estimation and surface thinning

As for curve reconstruction we consider a special point in a neighborhood with radius $H$, i.e. $\mathbf{x}_* \in \{\mathbf{x}_i\}$. This analysis is completely performed in real coordinates and not in voxel space. Using the same weighting function, Eq. (S1), we again perform weighted PCA to obtain the direction of minimal variance, corresponding to the normal direction of the plane which is locally tangential to the surface. Denoting the three eigenvalues of the scatter matrix $\lambda_1 \geq \lambda_2 \geq \lambda_3$, we can determine the linearity of the plane by considering the neighborhood parameter

$$\Lambda = \frac{\lambda_1 + \lambda_2}{\lambda_1 + \lambda_2 + \lambda_3} \in \left[\frac{2}{3}, 1\right]. \tag{S7}$$

The eigenvalues denote the rescaled (by the weights) variance of the point cloud in the eigendirections. For $\Lambda = 1$ we have a perfect plane and $\lambda_3 = 0$, while for $\Lambda = 2/3$ the point cloud has no preferred direction and is not planar. The neighborhood parameter thus serves as a measure of linearity analogous to the correlation in the curve thinning case.

To obtain a reasonable initial normal estimate, we optimize the neighborhood radius $H$ (for point $\mathbf{x}_*$, i.e. we optimize for every point in the point cloud) with respect to $\Lambda$. For this we start with $H_0 = 3\sqrt{(\Delta x)^2 + (\Delta y)^2 + (\Delta z)^2}$ for the lattice spacings in real space, increase by $\Delta H = \min\{\Delta x, \Delta y, \Delta z\}/2$ until either $\Lambda > 0.95$ or $H >= 5H_0$, taking the $H$ which maximizes $\Lambda$ in the latter case.

The obtained normals are oriented arbitrarily and need to be aligned. For this we propagate the normal from an arbitrary point across the surface along the EMST aligning the next normal on the previous one [9, 11].

As in curve thinning we now reconstruct the surface in the given neighborhood, replace $\mathbf{x}_*$ by the weighted centroid and project it onto a quadratic fit. This procedure again reduces the bias toward the direction in which the surface is bent, which occurs in a moving average (or K nearest neighbor) approach. Consider again $\mathbf{x}_* \in \{\mathbf{x}_i\}$ with the previously estimated normal $\mathbf{n}$ and a given radius $H$. We consider the differences $\mathbf{d}_i = \mathbf{x}_i - \mathbf{x}_*$, which are then expressed with respect to an orthonormal basis $(\mathbf{u}_x, \mathbf{u}_y, \mathbf{n})$, where we construct the $\mathbf{u}_{x/y}$ spanning the tangential space arbitrarily. We denote the components as $\hat{x}_i = \mathbf{d}_i \cdot \mathbf{u}_x$, $\hat{y}_i = \mathbf{d}_i \cdot \mathbf{u}_y$, and $\hat{z}_i = \mathbf{d}_i \cdot \mathbf{n}$. The weights are again given by Eq. (S1). We want to locally fit the function

$$f(x, y) = \frac{\beta_{xx}}{2} x^2 + \frac{\beta_{yy}}{2} y^2 + \beta_{xy} xy + \beta_x x + \beta_y y + \beta_0. \tag{S8}$$

For this we construct the design matrix

$$\mathbf{X} = \begin{pmatrix} \frac{\hat{x}_1^2}{2} & \frac{\hat{y}_1^2}{2} & \hat{x}_1 \hat{y}_1 & \hat{x}_1 & \hat{y}_1 & 1 \\ \vdots & \vdots & \vdots & \vdots & \vdots & \vdots \\ \frac{\hat{x}_{N_\text{sub}}^2}{2} & \frac{\hat{y}_{N_\text{sub}}^2}{2} & \hat{x}_{N_\text{sub}} \hat{y}_{N_\text{sub}} & \hat{x}_{N_\text{sub}} & \hat{y}_{N_\text{sub}} & 1 \end{pmatrix}, \tag{S9}$$

the weight matrix $\mathbf{W} = \text{diag}(w_1, \dots, w_{N_\text{sub}})$ and obtain the parameter vector $\hat{\beta} = (\beta_{xx}, \beta_{yy}, \beta_{xy}, \beta_x, \beta_y, \beta_0)$ through WLSE,

$$\hat{\beta} = \left(\mathbf{X}^T \mathbf{W} \mathbf{X}\right)^{-1} \mathbf{X}^T \mathbf{W} \mathbf{z}, \tag{S10}$$

with $\mathbf{z} = (z_1, \dots, z_{N_\text{sub}})$.

The initial assumption of a suitable $H$ is met if the neighborhood can be described well by the quadratic function in Eq. (S8). To assess this we compute the pseudo-coefficient of determination [10]

$$\mathcal{R}^2 = 1 - \frac{\left(\mathbf{z} - \mathbf{X}\hat{\beta}\right)^2}{\mathbf{z}^2}, \tag{S11}$$

which serves as a goodness of fit criterion and describes the proportion of explained variance in the unweighted reconstruction. Not including the weights in this formula accounts for the applicability the fit should have over a larger range to correctly describe the data: the fitted model should describe the data on the entire neighborhood of the observations without suppression via the weights [10]. In the denominator we do not consider the real mean for the variance but the variance from 0, contrary to Ref. [10], because the real mean may lie outside the surface and we are interested in the variance with respect to $\mathbf{x}_*$.

Taking $\mathcal{R}^2$ as a metric for the goodness of the quadratic fit, we optimize $H$: starting from $H_0$ as before, we increase by twice the $\Delta H$ as before, until either $\mathcal{R}^2 > 0.99$, or $H > 1.2 \max\left(\{\|\mathbf{x}_i - \mathbf{x}_j\| \mid i, j = 1, \dots, N_\text{sub}\}\right)$, or finally if the distance of select points from the paraboloid becomes too large, taking again the $H$ which maximizes our metric, $\mathcal{R}^2$, in the latter two cases. For the distance criterion we consider a cutoff distance of $d_\text{cutoff} = 2\sqrt{(\Delta x)^2 + (\Delta y)^2 + (\Delta z)^2}$ and compute the distance from the paraboloid via Newton iteration. This distance cutoff suppresses underfitting for very large $H$, where the reconstructed surface is outside the point cloud but still yields better $\mathcal{R}^2$.

To achieve surface thinning the special point is now projected onto the reconstructed surface, i.e. $\mathbf{x}_* \to \mathbf{x}_* + \beta_0 \mathbf{n}$.

#### Interfacial curvature calculation and normal correction

Curvature is calculated for the thinned surface. We again consider one specific point $\mathbf{x}_*$ of the thinned point cloud and perform a quadratic surface fitting with $\mathcal{R}^2$ optimization, as before. Now we do not perform any weighting, i.e. we choose the identity matrix $\mathbf{W} = \mathbf{I}$. We also consider a smaller cutoff distance of $d_{\text{cutoff}} = \sqrt{(\Delta x)^2 + (\Delta y)^2 + (\Delta z)^2}/4$ and an $\mathcal{R}^2$ threshold of 0.9998, using the same $H_0$ and $\Delta H$ as before, which we found to work well on our data.

Additionally, we prescribe $\beta_0 = 0$ and thereby enforce the quadratic surface to go exactly through our special point. For this the design matrix, Eq. (S9), is modified by deleting the last column, equivalent to neglecting the constant term in Eq. (S8). This suppresses potential underfitting at the edges of interfaces, caused by the one-sided distribution of points there. Using the inferred parameters, we correct the normal $\mathbf{n}$ by accounting for the linear component in the quadratic fit

$$\mathbf{n} \to \mathbf{n} - \frac{\beta_x}{1 + \beta_x^2 + \beta_y^2}\mathbf{u}_x - \frac{\beta_y}{1 + \beta_x^2 + \beta_y^2}\mathbf{u}_y. \tag{S12}$$

To calculate the curvatures corresponding to $\mathbf{x}_*$ we consider the (deflection) function describing our 2D surface locally, Eq. (S8), with the determined parameters (with $\beta_0 = 0$). The corresponding metric tensor describing the surface is [12]

$$\mathbf{g} = \begin{pmatrix} 1 + (\partial_x f)^2 & (\partial_x f)(\partial_y f) \\ (\partial_x f)(\partial_y f) & 1 + (\partial_x f)^2 \end{pmatrix}\Bigg|_{(x,y)=(0,0)} = \begin{pmatrix} 1 + \beta_x^2 & \beta_x \beta_y \\ \beta_x \beta_y & 1 + \beta_y^2 \end{pmatrix}. \tag{S13}$$

For the second fundamental form of the surface we have

$$\mathbf{h} = \frac{1}{\sqrt{\det \mathbf{g}}} \begin{pmatrix} \partial_{xx} f & \partial_{xy} f \\ \partial_{xy} f & \partial_{yy} f \end{pmatrix}\Bigg|_{(x,y)=(0,0)} = \frac{1}{\sqrt{1 + \beta_x^2 + \beta_y^2}} \begin{pmatrix} \beta_{xx} & \beta_{xy} \\ \beta_{xy} & \beta_{yy} \end{pmatrix}. \tag{S14}$$

The curvatures can now be computed through the Weingarten matrix

$$\mathbf{a} = \mathbf{h}\mathbf{n}^{-1}. \tag{S15}$$

If we denote with $c$ and $c'$ the principal curvatures then the mean curvature is $H = (c + c')/2$ and the Gaussian curvature $K = cc'$, which can be calculate from the Weingarten matrix via

$$H = \frac{1}{2}\operatorname{tr}\mathbf{a}, \qquad K = \det \mathbf{a}. \tag{S16}$$

As such we compute the mean and Gaussian curvature of the local quadratic fit for every point in the point cloud, giving access to a spatial distribution of curvatures along the surface.

#### Comparing reconstruction results to ground truth data

For a constant surface tension along an interface, we expect a constant mean curvature surface in the minimal energy configuration. The interfaces will thus resemble spherical segments quite often. To test our interface thinning algorithm, we consider a known surface which we then reconstruct. We consider a part from a spherical surface with radius $r = 15$ which we obtain for azimuthal angles $\phi \in [0.79\pi, 1.21\pi]$ and polar angles $\theta \in [0.29\pi, 0.71\pi]$. As for the curve reconstruction the interface is considered with a finite thickness $\delta r = 1$ and the corresponding voxels are determined via sampling. We then apply the surface thinning algorithm with the same hyperparameters as described before.

Supplementary Figure S8 depicts the thinning results for the voxelated surface. The original surface and the point cloud are shown in Supplementary Fig. S8a with a cut through the mid-plane $z = 0$, corresponding to the voxels $0 \leq z < 1$. The interface is subsampled with 4 points per voxel, just as in the case of real organoid interfaces. The thinned point cloud is projected well onto the real surface with only slight deviations, cf. Supplementary Fig. S8b. In Supplementary Fig. S8c the radial error is depicted, i.e. the distance of the points from the original surface, and we find a very good thinning result with an error within the $\pm 1/2$ voxel range. We find a slight overestimation of the radius, i.e. the points are projected onto a surface with a larger radius. This is, however, well within the uncertainty of the voxel segmentation. The error can be understood to result from the second-order Taylor approximation in the local fitting function. In order to minimize the quadratic deviation from the surface, a best fit will overestimate the quadratic contribution to compensate for the missing higher order terms, which leads to a negative constant term and thus a projection toward a larger radius.

We determined the normal angle error as the opening angle of the reconstructed normal at the thinned point and the corresponding theoretical normal at this point, cf. the histogram in Supplementary Fig. S8d. The errors are smaller then the angular errors for curve tangents and lie here in the range of $0^\circ$ to $4^\circ$. Note that this error might be larger for a few single points that sometimes cannot be projected well onto the real surface, depending on the subsampling points. This error is caused by enforcing the reconstruction in the curvature estimation to go through the points at which the reconstruction is performed. Albeit being larger at the surface edges, the normal angle error is well controlled. Considering the distribution across the thinned surface, cf. Supplementary Fig. S8b, the error seems to increase around voxel jumps due to the discontinuous nature of the original data.

Additionally to the overall smaller angular error the error at the surface corners seems to be even smaller, justifying the determination of tri-interfacial junctional tangents through the normal vectors of the corresponding interfaces.

For the reconstruction the mean curvature distribution is shown in Supplementary Fig. S8e. We see a broad distribution which peaks at a larger curvature than theoretically expected. The mean is slightly larger than we expect and lies just outside the interface thickness uncertainty $R \pm 1$ but is still comparable to that regime.

While the thinning algorithm overestimates the radius slightly the curvature reconstruction underestimates it.

#### Organoid geometry reconstruction

The presented surface thinning and curvature calculation algorithm has been optimized to reconstruct interfaces obtained via the 19-point-stencil in three dimensions. The premise of a three-dimensional point cloud which is obtained from volumetric data describing a finite-thickness surface is also given if we consider the epithelial monolayer as a finite-thickness material around the midplane, similar to modelling approaches [13, 14], and experimental analyses as elastic sheets [15–17]. As such the organoid segmentation itself, i.e. organoid versus background, is amenable to surface thinning as well. To reconstruct the organoid geometry we therefore employ the same surface algorithm as before, slightly adapting the hyperparameters of our approach. Consider the organoid thickness $t$ (i.e. the (local) average maximum distance of points in the organoid cells' volumes orthogonal to the mid-plane) which can be estimated from the microscopy data. We generate a point cloud from the organoid segmentation in a similar fashion as before, only that we now only subsample each voxel once and randomly retain approximately 25000 points in total, due to numerical limitations. The point cloud is again considered in real coordinates.

As already pointed out by Lee [8] for curve reconstruction, the algorithm can be applied iteratively to improve on the thinning result. The thickness of the organoid as a surface is much larger ($\approx 10 - 20\,\mu$m) than the one of cell-cell or cell-medium interfaces (a few voxels, corresponding to $\lesssim 1\,\mu$m) and as such we apply the algorithm for surface thinning twice. Compared to the interface reconstruction we do consider, however, changed values for the initial neighborhood radius $H_0$, radius stepping $\Delta H$, upper radius boundary $\hat{H}$ for the initial normal estimation, and the distance cutoff $d_{\text{cutoff}}$ for the different runs, as summarized in Supplementary Table S3.

To assess the quality of our organoid reconstruction, we construct a theoretical dumbbell-shaped organoid. For this we consider a rotationally symmetric organoid shell around the $x$ axis, which is described by its mid-plane. At the ends two half-spheres are considered with radii $R = 100$ voxels. The thickness of the half spheres is $\Delta R = 50$, i.e., we consider the volume with radius $R - \Delta R/2$ to $R + \Delta R/2$ from the spherical center. For the neck we consider a radial distance from the $x$-axis

$$r(\tilde{x}) = R - \frac{R}{3}\cos^2\left(\frac{\tilde{x}}{\tau\pi}\right), \tag{S17}$$

with center segment length $\ell$ and distance from the organoid's center $\tilde{x} \in [-\ell/2, \ell/2]$ along direction $x$. In the neck region, we consider the organoid to have a radial distance of $r - \Delta R/2$ to $r + \Delta R/2$. Note that this implies a varying thickness (in the normal direction of the mid-plane) along the symmetry axis.

For this shell volume the voxels which are inside the shell are considered. Contrary to single interface reconstructions the number of voxels in the entire organoid is so large that we need to reduce the number of points in the point cloud and not subsample. For both real organoids and the in-silico organoid considered here we only consider 25000 voxels, which are randomly selected from the volume and then subsampled with a single point.

Supplementary Figure S9 depicts the reconstruction of the in-silico generated test organoid. The sampled point cloud (Supplementary Fig. S9 a) is thinned and the mean curvature (Supplementary Fig. S9b) and Gaussian curvature (Supplementary Fig. S9c) are determined for the mid-plane on a point-to-point basis. Considering the points' radial distances from the symmetry axis of both the original point cloud (Supplementary Fig. S9d) and the thinned point cloud (Supplementary Fig. S9e) we find good thinning results toward the prescribed mid-plane, c.f. line in Supplementary Fig. S9e. The spatial error, i.e. the distance of the points from the prescribed mid-plane decreases substantially as the point cloud is thinned (Supplementary Fig. S9f). This can be seen when comparing the histograms of the spatial errors of the individual point clouds (Supplementary Fig. S9g).

The theoretical mid-plane curvatures along the $x$-axis are given by $H = 1/R$ and $K = 1/R^2$ in the spherical caps for mean curvature and Gaussian curvature, respectively. In the neck region the curvatures along the $x$-axis can be calculated using axial symmetry [12] to read

$$\begin{aligned} H(\tilde{x}) &= \frac{1}{2\,r(\tilde{x})\left(1+(r'(\tilde{x}))^2\right)^{1/2}} - \frac{r''(\tilde{x})}{2\left(1+(r'(\tilde{x}))^2\right)^{3/2}}, \\ K(\tilde{x}) &= -\frac{r''(\tilde{x})}{r(\tilde{x})\,(1+(r'(\tilde{x}))^2)^2}, \end{aligned} \tag{S18}$$

where the first and second derivatives of the radial mid-plane function $r$ with respect to $\tilde{x}$ are denoted as $r'$ and $r''$, respectively. To estimate the theoretical uncertainty of the curvature reconstruction, we consider the curvatures in the spherical cap regions for $R - \Delta R/2$ and $R + \Delta R/2$. In the neck region we consider $r - \Delta R/2$ and $r + \Delta R/2$ for the estimation. The reconstructed mean and Gaussian curvatures of the dumbbell are shown in Supplementary Fig. S9h,i, respectively. Our algorithm can reconstruct the organoid's mid-plane curvatures within the uncertainty from the organoid thickness. Note that the reconstruction smears out the discontinuities in the prescribed shapes' curvatures. This is to be expected due to the continuous quadratic fitting function assumed in curvature reconstruction. In real cellular systems, however, such discontinuities will be smeared out due to the finite size of individual cells anyways.

### Supplementary Note 2: Force and pressure inference method

For force inference we assume that the morphology can be described completely by a tension-based approach and that for each interface separating two cells the surface tension is constant. For junctions (cf. Fig. 1c) the possible line tension (or tangential force component at the two ends) is also assumed to be constant.

#### Inferring interfacial tensions from dihedral angles

Based on our three-dimensional reconstruction methods (see Supplementary Note 1) we have thinned junctional and interfacial points with corresponding normal vectors for the interfaces.

To infer interfacial tensions, which serve as a proxy for contractile forces, we assume mechanical equilibrium at the tri-interfacial junctions at which three interfaces meet. In the normal space of the junction the interfaces exert a force on the junction from surface tension. The relative magnitude of these forces influences the dihedral angles between the interfacial tangential spaces, cf. Supplementary Fig. S12a. To reconstruct the dihedral angles, we start off from the normal vectors of the interfaces at the junctional point. For this we consider the normal vectors $\mathbf{n}_i$ of the (thinned) interfacial point closest to the (thinned) junction point. These normals should now be co-planar and span the normal space of the junction, i.e. the space orthogonal to the junctional tangent $\mathbf{j}$. In reality this is not necessarily the case and we determine the averaged (normalized) junctional tangent as [11]

$$\hat{\mathbf{j}} = \frac{\mathbf{n}_1 \times \mathbf{n}_2 + \mathbf{n}_2 \times \mathbf{n}_3 + \mathbf{n}_3 \times \mathbf{n}_1}{\|\mathbf{n}_1 \times \mathbf{n}_2 + \mathbf{n}_2 \times \mathbf{n}_3 + \mathbf{n}_3 \times \mathbf{n}_1\|}. \tag{S19}$$

The normal vectors are then projected into the orthogonal space to obtain the co-planar normals $\hat{\mathbf{n}}_i$, cf. Supplementary Fig. S12b.

Force balance in the junctional normal space now implies

$$\begin{aligned} 0 &= \gamma_1 + \gamma_2 \cos\theta_{12} + \gamma_3 \cos\theta_{13}, \\ 0 &= \gamma_2 \sin\theta_{12} - \gamma_3 \sin\theta_{13}, \end{aligned} \tag{S20}$$

with dihedral angles $\theta_{ij}$ and interfacial tensions $\gamma_i$. We selected an arbitrary interface as reference to choose our coordinate system. Note that force balance can be written down in different forms [18], but the form used here yields good reconstruction results. In the junctional normal space the interfacial normals and tangents are now orthogonal and the dihedral angles are computed, cf. Supplementary Fig. S12c.

Equations (S20) are linear in the interfacial tensions $\gamma_i$ and for every junction we obtain two such equations. Introducing the tension vector $\boldsymbol{\gamma} = (\gamma_1, \ldots, \gamma_{N_{\mathrm{int}}})$ for interfaces $i = 1, \ldots, N_{\mathrm{int}}$, we may write this as a matrix equation

$$\mathbf{M}_\gamma \boldsymbol{\gamma} = \mathbf{b}_\gamma. \tag{S21}$$

The system of equations determines the relative tension magnitudes but does not contain information on the absolute value. As such we augment the system with an additional equation

$$\sum_{i=1}^{N_{\text{int}}} \gamma_i = N_{\text{int}}, \tag{S22}$$

i.e. the average tension is set to 1. All these equations are now summarized into the design matrix $\mathbf{M}_\gamma$ with residual vector $\mathbf{b}_\gamma = (0, \ldots, 0, N_{\text{int}})$. In general, the dihedral angles may vary along the junction. We consider the averaged dihedral angles for tension inference. In the organoid experiments we find this to be realized, contrary to what has been observed in embryos [11].

The resulting system of equations is overdetermined and the matrix equation cannot be inverted exactly. Knowing all the dihedral angles defining $\mathbf{M}$, we may therefore infer the tension vector via least squared error estimation to be

$$\boldsymbol{\gamma} = \left(\mathbf{M}_\gamma^T \mathbf{M}_\gamma\right)^{-1} \mathbf{M}_\gamma^T \mathbf{b}_\gamma. \tag{S23}$$

#### Pressure inference from interface mean curvatures

Assuming the interface shape is driven by surface tension, the Young-Laplace equation holds across the interfaces. Therefore a pressure difference between cells $i$ and $j$, $\Delta P = P_i - P_j$, leads to a constant mean curvature interface between the cells with mean curvature $H_k$ and surface tension $\gamma_k$. The equation reads

$$P_i - P_j = 2H_k\gamma_k, \tag{S24}$$

which is linear in pressures, cf. Extended Data Fig. 4a. The interfacial tensions are determined via inference and the mean curvatures are known for the (thinned) interfacial points. Considering the averaged mean curvatures across the interface we obtain the matrix equation

$$\mathbf{M}_{\text{p}}\mathbf{p} = \mathbf{b}_{\text{p}}, \tag{S25}$$

with pressure vector $\mathbf{p} = (P_1, \ldots, P_{N_{\text{c}}})$ for cells $i = 1, \ldots, N_{\text{c}}$ and residual vector $\mathbf{b}_{\text{p}} = (2H_1\gamma_1, \ldots, 2H_{N_{\text{jun}}}\gamma_{N_{\text{jun}}}, 0)$ for junctions $i = 1, \ldots, N_{\text{jun}}$. The augmented design matrix $\mathbf{M}_{\text{p}}$ contains Eqs. (S24) for each junction and the additional condition $P_j = 0$ for the outside (background) pressure. Since the Young-Laplace relation only describes pressure differences, pressures are only determined up to an additive constant. We therefore consider the pressure difference with the background. In addition we do not know the absolute scale of the surface tensions and as such can only calculate pressures per average tension.

The pressures are again inferred via least squared error estimation

$$\mathbf{p} = \left(\mathbf{M}_{\text{p}}^T \mathbf{M}_{\text{p}}\right)^{-1} \mathbf{M}_{\text{p}}^T \mathbf{b}_{\text{p}}. \tag{S26}$$

Pressures and surface tensions can be inferred simultaneously [11], by writing the two matrix equations as one, or the pressure can be inferred after the tensions. We perform the inferences separately as the curvature estimation and pressure errors should not feed into the inference of the interfacial tensions.

#### Junctional tangential force inference

In the inference of interfacial surface tensions we have considered force balance in the normal space of the tri-interfacial junctions. Assuming mechanical equilibrium, possibly existing forces in the tangential space must also be balanced, which implies force balance at points where multiple junctions meet, cf. Supplementary Fig. S13a. These points occur at the apical and basal ends of junctions – we have called them junctional vertices. Assuming that the force magnitude is identical at both junctional ends (as would be the case for elastic springs or line tensions), force balance at the junctions reads

$$\sum_{i=1}^{4} \tau_i \mathbf{j}_i = 0, \tag{S27}$$

with line force magnitudes $\tau_i$ and normalized junctional tangents $\mathbf{j}_i$ for the junctions involved in the vertex $i = 1, \ldots, 4$ (in this case 4), cf. Supplementary Fig. S13b.

For each junctional vertex we have a force balance equation, Eq. (S27), which is linear in the tangential forces. Introducing the tangential force vector $\boldsymbol{\tau} = (\tau_1, \ldots, \tau_{N_{\text{vtx}}})$, this again yields a linear matrix equation

$$\mathbf{M}_\tau \boldsymbol{\tau} = \mathbf{b}_\tau, \tag{S28}$$

with the augmented design matrix $\mathbf{M}_\tau$, containing the force balance equations, and the residual $\mathbf{b}_\tau = (0, \ldots, 0, N_{\text{vtx}})$. As for surface tensions, the tangential forces are only known with relative magnitudes and we set the average value to 1 by introducing the equation $\sum_{i=1}^{N_{\text{vtx}}} \tau_i = N_{\text{vtx}}$. This equation is again solved in a least squared sense via

$$\boldsymbol{\tau} = \left(\mathbf{M}_\tau^T \mathbf{M}_\tau\right)^{-1} \mathbf{M}_\tau^T \mathbf{b}_\tau. \tag{S29}$$

#### Regularization of the inverse problem

All aforementioned inference problems in force and pressure inference are inverse linear problems. Since these problems are overdetermined and inherently noisy, a least squared error approach has been suggested when inverting the matrix equation by using the pseudoinverse

$$(\mathbf{M}_x^T \mathbf{M}_x)^{-1} \mathbf{M}_x^T \tag{S30}$$

for $x = \gamma, p, \tau$. The resulting estimator for $\mathbf{x}$ then minimizes the least squared error

$$\|\mathbf{M}_x \mathbf{x} - \mathbf{b}_x\|^2. \tag{S31}$$

Due to errors from incorrect segmentations and badly reconstructed angles and curvatures, the problem may become ill-posed and very sensitive to noise, leading to an ill-conditioned matrix $\mathbf{M}_x^T \mathbf{M}_x$, which cannot be inverted properly. This problem also occurs in different inverse problems such as the traction estimation from deformations in traction force microscopy [19, 20].

To resolve this, Thikonov regularization is applied, i.e. the regularized least squares error

$$\|\mathbf{M}_x \mathbf{x} - \mathbf{b}_x\|^2 + \lambda \|\mathbf{x}\|^2 \tag{S32}$$

is minimized. The minimizer, which serves as an estimator for the quantity to infer, $\mathbf{x}$, then reads

$$\mathbf{x} = (\mathbf{M}_x^T \mathbf{M}_x + \lambda \mathbf{I})^{-1} \mathbf{M}_x^T \mathbf{b}_{\mathrm{x}}, \tag{S33}$$

with identity matrix $\mathbf{I}$. This regularization suppresses high noise-driven fluctuations and is more stable with respect to erroneous data, as we expect it from wrongful segmentations. The parameter $\lambda$ is chosen such that it minimizes Allen's Predicted Residual Error Sum of Squares (PRESS) statistics, which is an optimality criterion in leave-one-out cross validation [21]. The optimal $\lambda$ is efficiently calculated minimizing the PRESS statistics numerically for the data at hand [22].

### Supplementary Note 3: In-silico verifications of the force inference and pressure method in the two-cell and bubbly vertex model system, and for peaked distributions

We test the full inference method using synthetic in-silico data. For such data inference results can be compared against the underlying ground truth data that was used during generation of the synthetic data.

#### Verification of force and pressure inference in the two-cell system

We first consider a system of two cells, sharing a spherical cap interface, where we can easily calculate pressures and tensions based on the shapes. Inferring surface tensions of two spherically shaped cells or vesicles [23, 24] is a prototypical case of force inference due to its simplicity. As such it has been proposed before as a test case to benchmark three-dimensional force inference approaches [11].

To verify force inference capabilities of our algorithm, artificially created segmentation data for such a spherical two-cell system with prescribed Radii $R_1$ and $R_2$ of the two cells, respectively, and a spherical cap boundary $R_3$ is analyzed. A sketch of the situation with the corresponding created segmentation and the reconstruction and inference results are shown in Extended Data Fig. 1. Note that due to the small size of the linear system no Thikonov regularization has been used here, as this would lead to over-regularization and suppression of the real tension deviations: The single junction and the setting of the average tension yield three equations for the three unknowns, making the system non-over-determined and well-posed. We find that our reconstruction algorithm can reconstruct the three individual interfaces and the circular junction well from the data. At each point along the junction we obtain tangent vectors for the individual membranes (cf. Extended Data Fig. 1e), which allow the determination of the dihedral angles along the junction contour (cf. Extended Data Fig. 1f). We see that the angles are constant around the junction with fluctuations stemming from two possible sources of error: (i) the reconstruction will produce statistical uncertainties, yielding noise, and (ii) the membrane normals are determined from voxelated data and the normals are taken from membrane points around the junctional positions, where the point cloud data also comes from the voxelated segmentations. In the original point cloud representation of the junction (cf. Extended Data Fig. 1d) we see that the points are from two neighboring voxel sheets, explaining the discrete nature of the angle fluctuations. Comparison to the ground-truth angles (cf. Extended Data Fig. 1g) shows that our algorithm reconstructs the angles via averaging.

As has been noted before [11, 25], the reconstruction error strongly depends on the underlying image resolution. To quantify this, we performed force inference on different sizes of the two-cell system for different curvatures of the separating interface (cf Extended Data Fig. 1h) and calculated the total angular error (TAE)

$$(\text{TAE}) = \sum_{\{i,j\}} \left| \hat{\theta}_{ij} - \theta_{ij} \right| , \tag{S34}$$

with summation over all angles and with inferred and real angles $\hat{\theta}_{ij}$ and $\theta_{ij}$, respectively. The error is of the same order of magnitude as what can be expected from the voxelization of the interfaces and thus the uncertainty in the exact radii. It generally decreases with higher resolution but shows a strong dependence on the relative size of the cells and of the exact resolution, which results in offsets on the voxelated grid and thus fluctuating noise in the angles. As we set the average surface tension to 1, the relative error of the deviation from this average value is of interest, i.e. the relative tension deviation error (RTDE)

$$(\text{RTDE}) = \left\langle \left| \frac{\hat{\gamma}_i - \gamma_i}{\gamma_i - 1} \right| \right\rangle , \tag{S35}$$

with inferred and real surface tensions $\hat{\gamma}_i$ and $\gamma_i$, respectively. This error also decreases with resolution, cf. Extended Data Fig. 1i, and we find that for the different relative cell sizes we always find errors in the range of the expected error from voxel resolution. The resolution in experimental images is of the order of a total membrane diameter (on apical and basal side) of up to 30-40 voxels (in experimental data this corresponds to an interface size of $7.5 - 10\,\mu$m), indicating that the error will at least be in the range of 5% of the average tension.

To investigate the accuracy of the pressure inference, in Supplementary Fig. S14 we investigated a two-cell system with identical surface tensions for the three interfaces but with prescribed cellular pressures. According to the Young-Laplace relation, Eq. (S24), this then leads to three different spherical cap radii, which were reconstructed via curvature inference. The average relative curvature error, cf. Supplementary Fig. S14e, is of the order of 4%. The inferred curvatures are not fully spatially homogeneous even though they should be (cf. Supplementary Fig. S14a,b), and we systematically overestimate them (cf. Supplementary Fig. S14d). This can be explained by the fitting procedure. Despite the weighting in the paraboloid fit and the cut-off in maximum distance, the quadratic Taylor expansion is of interest, which acts as an osculating surface and not an error-minimizing surface. The alternative approach of directly fitting constant mean curvature surfaces (spherical caps, cylinder segments, and planes) has not yielded more satisfactory results either.

The resulting relative pressure error is also in the range of 5%, cf. Supplementary Fig. S14f, and of comparable size to the tension error. Note that due to the occurrence of both tension and pressure in the Young-Laplace equation, Eq. (S24), the pressure reconstruction will be limited by the tension error anyways.

#### Verification of pressure inference in the bubbly vertex model

After verification on the two-cell test case, the pressure inference was also tested on an in-silico organoids simulated with the bubbly vertex model. Interface averages of reconstructed curvatures display good agreement with ground truth simulated in-silico data with a slight overestimation of large mean curvatures (Supplementary Fig. S15a). This allows us to infer both cellular and luminal pressures, where cellular pressures show a strongly peaked distribution due to the minimal pressure differences between the cells with a few outliers (Supplementary Fig. S15b). Luminal pressure can be modulated in-silico by imposing a volume constraint on the lumen. For different luminal volumes we found varying inferred luminal pressures, which qualitatively match the theoretical values with a constant offset (Supplementary Fig. S15c). This is a result of the overestimation of large curvatures by the inference algorithm. Similar to the tensions, we cannot reconstruct absolute pressure values, and therefore determine pressures per average tensions. Thus, we cannot fully infer the absolute luminal and cellular pressures but our method robustly reconstructs differences in pressures (per average tension) between individual samples.

##### Reconstruction limits for peaked multimodal distributions

In order to test the limits of the force inference method we next considered a budded organoid with very strong patterning compared to the variance inside the individual regions of the budded organoid. This is much closer to the theoretically often-used mean-field approach [26, 27].

Supplementary Figure S16 depicts the force inference results for this simulation. We see that the reconstructed distributions (Supplementary Fig. S16b,e) are not as separated and peaked as the ground truth data (Supplementary Fig. S16a,d). The multimodality in the total distribution cannot be reconstructed and the relative tension difference between the groups not perfectly resolved. Looking at the correlation of apical tensions, we find that the large apical tension in the buds cannot be reconstructed (Supplementary Fig. S16c). However, our algorithm is able to correctly reconstruct the lower apical tension in the neck region. To rule out that this shortcoming is an effect from Thikonov regularization, which is equivalent to the assumption of Gaussian distributed parameters that are to be inferred, we also considered a reconstruction without regularization (Supplementary Fig. S16f). The same qualitative disagreement of the apical tensions is found but the noise in the reconstruction is severely increased.

One possible reason for the error-prone inference result is the much smaller apical interfaces in the buds due to the strong curvature, increasing the reconstruction uncertainty. Moreover, the statistical approach is likely unable to perfectly resolve distributions which are so strongly peaked and add up to a distribution of all tensions that is not unimodal. However, in biological samples a larger variance between the different tensions is to be expected, justifying the regularization procedure.
